# Conversion of marginal land into switchgrass conditionally accrues soil carbon and reduces methane consumption

**DOI:** 10.1101/2020.03.18.997304

**Authors:** Colin T. Bates, Arthur Escalas, Jialiang Kuang, Lauren Hale, Yuan Wang, Don Herman, Erin E. Nuccio, Xiaoling Wang, Ying Fu, Renmao Tian, Gangsheng Wang, Daliang Ning, Yunfeng Yang, Liyou Wu, Jennifer Pett-Ridge, Malay Saha, Kelly Craven, Mary Firestone, Jizhong Zhou

## Abstract

Switchgrass (*Panicum virgatum* L.) is a perennial C_4_ grass native to tallgrass prairies of the Central US, and a promising bioenergy feedstock. Switchgrass can be cultivated on soils with low nutrient contents and its rooting depth, of up to 2 m, has brought attention to the crop as a potential mechanism to sequester and build soil carbon (C). Switchgrass, therefore, offers multifaceted benefits on degraded soils by enhancing soil organic matter content. However, to evaluate the sustainability of switchgrass-based biofuel production, it is crucial to understand the impacts of land conversion and switchgrass establishment on biotic/abiotic characteristics of various soils. In this study, we characterized the ecosystem-scale consequences of switchgrass growing at two highly-eroded, ‘Dust Bowl’ remnant field sites from Oklahoma US, with silt-loam (SL) or clay-loam (CL) soil textures having low nitrogen (N), phosphorus (P), and C contents. Paired plots at each site, including fallow control and switchgrass-cultivated, were assessed. Our results indicated that switchgrass significantly increased soil C at the SL site and reduced microbial diversity at the CL site. The CL site exhibited significantly higher CO_2_ flux and higher respiration from switchgrass plots. Strikingly, switchgrass significantly reduced the CH_4_ consumption by an estimated 39% for the SL site and 47% for the CL site. Structural equation modeling identified soil temperature, P content, and soil moisture levels as the most influential factors regulating both CO_2_ and CH_4_ fluxes. CO_2_ flux was also influenced by microbial biomass while CH_4_ flux was influenced by microbial diversity. Together, our results suggest that site selection by soil type is a crucial factor in improving soil C stocks and mitigating greenhouse gas (GHG) fluxes, especially considering our finding that switchgrass reduced methane consumption, implying that carbon balance considerations should be accounted for to fully evaluate the sustainability of switchgrass cultivation.

## Introduction

Taking place over three waves during the 1930s, The American ‘Dust Bowl’ was a catastrophic ecological disaster that brought severe drought and dust storms to the central prairies and affected roughly 40 million hectares of land (Schubert et al., 2004; Worst, 1982; Baumhardt, 2003). Strong wind erosion exacerbated the topsoil displacement, creating many ‘marginal’ lands of low soil nutrient quality across Oklahoma and the Southwestern US (Texas, Kansas, Colorado, and New Mexico). Since then, many of these sites have remained a ‘poor’ fit for agricultural development. It has been suggested that the implementation of deep-rooted perennial grasses may aid in soil restoration at these sites, offering economic benefits to farmers in the form of cellulosic feedstock for bioenergy production (Gelfand et al., 2013).

Switchgrass (*Panicum virgatum* L.), a tall perennial deep-rooted grass native to the Central North American Plains, is an auspicious bioenergy crop suitable for future large-scale cultivation in the US (McLaughlin and Kszos, 2005). This enthusiasm for switchgrass stems from its excellent ability to exhibit high biomass even on low-quality soils unfit for traditional agricultural practices, with little to no additional inputs (Tilman et al., 2006). Long-term cultivation experiments have suggested that switchgrass can improve soil productivity through the net input of soil C (Gelfand et al., 2013, Ma et al., 2000). Therefore, large-scale switchgrass cultivation may offset GHG emissions and serve as a means for improved soil fertility through C sequestration at nutrient-poor sites (Anderson-Teixeria et al., 2009). Switchgrass is also known to be highly drought tolerant (Barney et al., 2009) and can prevent topsoil erosion-two major problems in the Central and Southwestern USA including states of OK, TX, KS, CO, and NM. It is estimated that 15 million hectares of arable land would need to be converted into biofuel crop in order to meet the US Department of Energy’s plan to replace 30% of transportation fossil fuels with biofuels by 2030 (Bouton, 2007; Bouton, 2008). An estimated 11% of the contiguous US is considered nutrient-poor or ‘marginal’ land (Milbrandt et al., 2014) and currently represent an under-utilized resource that may be well suited for switchgrass cultivation (Stoof et al., 2015). Since perennial crop systems have high root biomass and exudates, they can improve soil C stability and aggregate formation (Tiemann and Grandy, 2015). Like other perennial crops, Switchgrass has been broadly associated with increases in soil C at many different sites in the central and northern Great American Plains (Liebig et at., 2008; Zan et al., 2001; Frank et al., 2004; Dabney et al., 2004). However, this potential C accrual may be offset by higher soil respiration arising from stimulated microbial C mineralization or increased root respiration.

Because soil microorganisms are critical drivers of soil nutrient cycling, understanding plant-microbe interactions during switchgrass cultivation could inform land management strategies toward the promoting of soil nutrient acquisition and recycling, along with reducing GHG emissions. By examining microbial ecology of switchgrass influenced systems, researchers have revealed mechanistic understanding of the ways it enhances ecosystem services such as C sequestration, soil fertility, and GHG emissions (McLaughlin and Kszos, 2005; Ker et al., 2014; Clarck et al., 2005; Bahulikar et al., 2014; Ghirmire et al., 2009; Kim et al., 2012; Ghimire and Craven, 2011). For instance, it was shown that N fertilization, at least at some sites, did not increase soil-surface carbon dioxide (CO_2_) emissions despite increases in above (Mulkey et al., 2006; Lee at al., 2007) and below ground biomass (Sher et al., 2020). However, the relative impact of methane (CH_4_) emissions during land conversions are not yet fully understood (Monti et al., 2012; Robertson and Grace, 2004; Fritsche et al., 2010). Additionally, the ecological consequences of land conversion, its impact on soil microbial ecology and functionality, as well as the overall sustainability of switchgrass cultivation as a biofuel crop remain to be demonstrated. Further, only a few studies have evaluated switchgrass cultivation at sites with low soil N, C, or P contents or in marginal lands that have experienced high rates of topsoil erosion (Gelfand et al., 2013; Ashiq et al., 2017). Particularly, we currently have a very limited understanding of how the transition from mixed annual grassland communities to switchgrass row-crop systems can affect (i) soil geochemical composition at nutrient-poor sites; (ii) soil microbial biodiversity, and (iii) the overall prairie ecosystem functionality, specifically GHG fluxes.

In this study, we monitored the ecosystem-level effects of switchgrass establishment over two consecutive growing seasons (n = 17 months) in two nutrient-poor (relatively low N, P, and C contents) field sites in Southern Oklahoma (designated SL for the silt-loam soil texture and CL for the clay-loam soil texture). We compared switchgrass and natural fallow plots in terms of soil chemistry (C, N, and P), soil GHG fluxes (CO_2_, CH_4_, and N_2_O), and microbial community composition. We tested the hypotheses that annual mixed grassland conversion to switchgrass (i) increases topsoil C levels over time; (ii) switchgrass increases CO_2_ respiration while maintaining similar CH_4_ emission and N_2_O flux relative to annual mixed grassland communities (fallows); and (iii) switchgrass modifies the microbial community during establishment and decreases species richness over time. We expect shifts in microbial community composition to correlate with observed GHG fluxes. Our results indicated that switchgrass had a site-specific, with increased CO_2_ respiration and decreases in microbial species richness observed only at the CL site, while soil C accumulation was observed only for the SL site. Switchgrass significantly reduced the methane consumption rates regardless of soil type.

## Methods

### Field site, soil sampling, and root biomass estimation

Samples were taken from two sites in southern Oklahoma, a silt-loam site (SL) near the Texas border (34.18691°N, −97.08487°W) and a clay-loam site (CL) in Ardmore (34.172100°N, −97.07953°W). Prior to our experiment, each field site had experienced annual crop rotation and periods of being left fallow. At each site, a switchgrass field plot (27 x 22 m) containing 500 genetically distinct individuals of the lowland Alamo variety with a 1 m spacing between plants and a corresponding fallow plot (27 x 22 m) were established in the summer of 2016 (Fig. S1a,b). All plots were tilled before planting switchgrass. Fallow plots were allowed to undergo natural succession of grasses and weeds over the time course of the experiment. To allow GHG measurements, at each plot, 21 PVC collars (diameter 23.63 cm x 12.8 cm x 1 cm) (Fig. S1c) were embedded 8 cm into the soil and placed in a cross design (Fig. S1a,b) with five collars extending in each cardinal direction from a central origin collar at the center of each plot. After trace gas measurement from each collar, two soil cores (15 - 20 cm in depth) were taken from within a 20 cm radius of each collar (Fig. S1d), thoroughly mixed, and separated into two bags, one for geochemical analyses and one for DNA extraction. Sampling flags were placed to prevent re-sampling the same location twice and each core was filled by topsoil taken from outside the plot. All soil samples were immediately stored on ice, transported back to the lab and kept at either 5□ for geochemical analyses or −80□ for DNA extraction. In May 2017, total belowground root biomass was estimated using the Fraiser et al., 2016 method. Briefly, four 0-1 m cores were taken between six target plants and an adjacent plant (Fig. S2b). Soil cores were then divided into 5 depths for every 20 cm of soil. For each depth, roots were extracted from cores through sieving and soaking the soil in water. Roots from each layer of soil were collected, dried and massed from each layer. Switchgrass root biomass was estimated as described elsewhere (Frasier et al., 2016). For fallow plots, four randomly assigned 1 m^2^ subplots within each of the quadrants were selected to take four 0-1 m soil cores to represent root biomass across the plot (Fig. S2c). No roots were detected from fallow soil cores below 60 cm depth.

### Soil geochemistry, pH, and moisture

Soil pH, moisture, total available C, N, P, nitrate (NO_3_), and ammonium (NH_4_) were measured following as described elsewhere (Hendershot et al., 2007). Briefly, 10 g of soil was placed into a 50 ml tube with distilled H_2_O added to the 50 ml fill line. Tubes were gently shaken for 30 minutes and given an hour to settle before pH measurement using a pH probe (Acccumet excel XL15 pH meter, Fisher Scientific, Hampton NH, USA). Soil moisture was determined by a gravimetric drying protocol that dried > 5 g of soil for one week at > 60 □ before re-massing to establish the percent of water lost after drying. To determine other soil geochemical parameters, soil samples were dried in an oven at 60 □ for a week followed by sieving to remove unwanted material with a 4 mm sieve. Soil samples were then shipped seasonally to the Oklahoma State University (OSU) soil testing lab where Mehlich III extractions (to quantify the available P in the soil), KCL extractions (to determine NH_4_ and NO_3_ concentrations) were performed and total soil C/N amounts were measured via dry combustion (LECO corporation, St. Joseph MI, USA).

### Environmental parameters

Daily environmental data for 21 different environmental variables (at 5 to 15-minute resolution) were obtained from two weather monitoring stations (Ardmore and Burneyville) belonging to the Oklahoma MESONET network (http://mesonet.org/) that were the closest to our field sites (1.43 km and 2.3 km from SL and CL, respectively). Variables used included air temperature, bare soil temperature, covered soil temperature, atmospheric pressure, relative humidity, and precipitation (Table S1 and Table S2).

### Trace gas fluxes

The CO_2_, CH_4_, and N_2_O fluxes were measured monthly via cavity ring-down spectrometry using a Picarro G2508 analyzer (Picarro, Santa Clara, CA, U.S.A.). Measurements were taken continuously every 2 seconds from a total of 6 minutes per collar. This allowed us to obtain gas concentrations in parts per million. Raw data from each gas were separated and then manually inspected to remove the beginning and the end of the measurements, which are often influenced by the pushing/pulling of the gas chamber. Then three models (linear, quadratic, and exponential) were fitted for each sample and gas species to characterize the variation of gas concentrations across time and the ‘best model’ was selected based on AIC scores. Flux estimations for each of the gases were then calculated using the following equation (Christiansen et al., 2015):

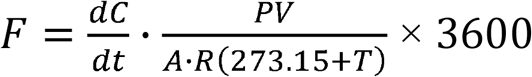

Where 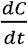 is the slope of the best fitted model at t = 0, V is the chamber volume (L), A is the chamber area (m^2^), R is the gas constant in L atm K^−1^ mol^−1^, T is the temperature in Celsius, when the chamber pressure is assumed to be equal to 1 atm. The 3600 factor is included to convert the flux to hourly values. For CO_2_ flux, F was then divided by 1000 to obtain the correct units of millimoles per m^2^ per hour.

### Soil DNA extractions, microbial community sequencing and analysis

A freeze grinding method (Zhou et al., 1996) was combined with the Powersoil DNA extraction kit (Qiagen, Venlo, Netherlands) to extract DNA from a total of 1,428 soil samples, which typically yielded soil DNA of both high quantity and quality. For microbial community profiling a two-step PCR method (Wu et al., 2015) was used for amplification of the V4 region of the bacterial 16S rRNA gene using the 515F, 5’-GTGCCAGCMGCCGCGGTAA-3’ and 806R, 5’-GGACTACHVGGGTWTCTAAT-3’ primers. Sequencing of the 16S rRNA gene amplicons was conducted on the Illumina Mi-Seq DNA sequencing platform (Illumina Inc., San Diego, CA, U.S.A.). Amplicon sequence data was analyzed using an internal pipeline known as the Amplicon Sequencing Analysis Pipeline (Zhang et al., 2014) (ASAP, version 1.4). MiSeq sequences were quality checked with FastQC (version 0.11.5), pair-end sequences were merged based on their 3’ overlap using PEAR (version 0.9.10) with a quality score cutoff set to 20, and assembly length between 200-400 with the minimum overlap length set to 50 bp. The program *split libraries_fastq.py* from the QIIME package (Kuczynski et al., 2012) (version 1.9.1) was used to assign reads to each sample (demultiplexing) based on the barcodes for each individual sample with a maximum allowed barcode error of 0 and the trimming quality score set to 20. Primer sequences were then trimmed and removed. Sequences from multiple split libraries were merged together. Dereplication was performed by USEARCH (Edgar, 2010) (version 9.2.64) using the command *fastx_uniques* (utilizing the size-out option for sequence abundance output). Operational Taxonomic Units (OTUs) were clustered using UPARSE, with the OTU identity threshold set to 0.97 and the singletons/chimeric sequences removed (Edgar, 2013). The OTU table was generated by the command *-usearch_global* in USEARCH. Each representative sequence for each OTU was classified with the RDP Classifier (Wang et al., 2007) (16S: training set 16, June 2016) with the confidence cutoff set to 0.8. OTUs in the 16S sequence reads assigned to Chloroplast at the Order level were removed. Representative sequences for each OTU were used to construct a phylogenetic tree. Sequences were then aligned using MAFFT (Katoh, 2002) (version 3.8.31) and alignments were filtered using Gblocks (Castresana, 2000) (version 0.91b) with the options -t=d, -b4=3 and -b5=h. FastTree (Price et al., 2009) was used for constructing the phylogenetic tree using the filtered alignments. The phylogenetic tree and OTU tables were used to calculate alpha diversity (phylogenetic based indexes) and beta diversity (UniFrac distance) using programs packaged in QIIME (Caporaso et al., 2010) and R.

### Statistical analyses

All analyses were conducted using R statistical software (3.4.4, R Core Team, 2014) and figures were produced using the package ggplot2 (Wickham, 2009). Data normality was tested using the Shapiro test. We tested for differences between plots in GHG flux and microbial alpha diversity by using linear mixed models to correct for repeated measurements (i.e. collars within plots) and to analyze the data over time (R package *lme4*, Bates et al., 2015). Pairwise comparisons for soil respiration between treatments were conducted using Wilcoxon Rank Sum test and effect sizes were calculated using Mann-Whitney U Test. Differences in soil biogeochemical properties between treatment were tested using Kruskal-Wallis test and effect size was calculated using epsilon squared. Soil geochemical dissimilarity was calculated from scaled data using Euclidean distances (*vegan* R package). Then mean dissimilarity across collars was used to construct linear mixed models to view changes in dissimilarity over time. Differences in microbial community structure across plot, site and time were tested using PERMANOVA test based on Bray Curtis and weighted-UniFrac dissimilarity for taxonomic and phylogenetic diversity, respectively. Differences in relative abundance between groups and time points was calculated by multiple Student T-Tests and p-values were adjusted by conservative Bonferroni correction to compensate for increased Type 1 errors over multiple time points.

Structural equation modeling (SEM) were used to explore the direct and indirect relationships among environmental variables and GHG fluxes (CO_2_ and CH_4_) at either site. We first considered a full model that included all reasonable pathways, then eliminated nonsignificant pathways until we obtained a final model with only significant pathways. We used a *χ*^2^ test and the root mean square error (RMSE) to evaluate the fit of our model. The SEM-related analysis was performed using the lavaan R package (Rosseel, 2012).

## Results

### Changes in soil geochemistry

The conversion of grassland into switchgrass appeared to have a site-specific impact on soil geochemistry. A principal component analysis (PCA) of the soil geochemistry data revealed strong differences between the two sites (Figure 1). Heterogeneity of the geochemical parameters was higher at the CL site, as displayed by the dispersion of blue samples in Figure 1.

**Figure 1.**
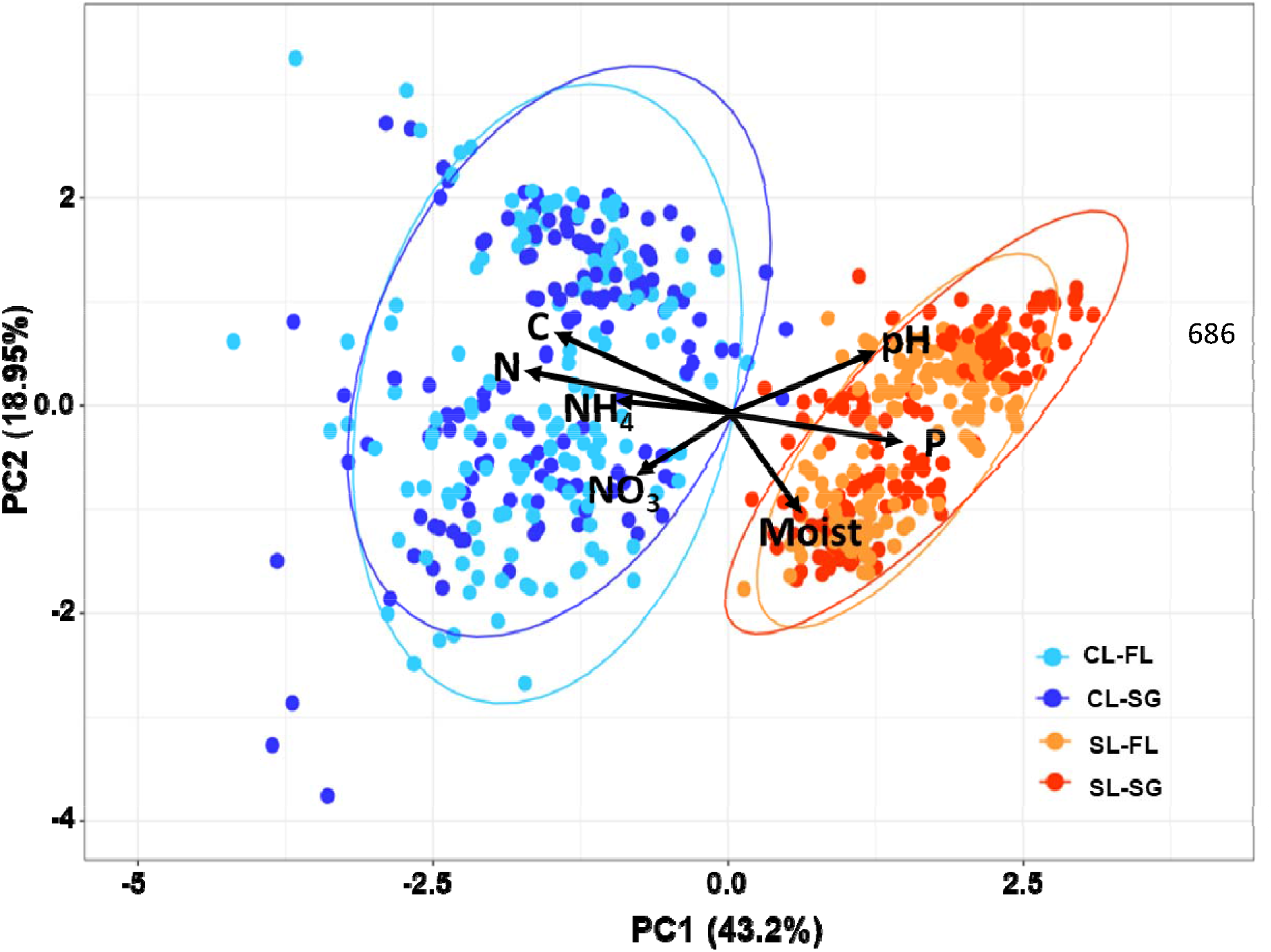
Differences in soil geochemical properties between the two studied sites. Principal component analysis. Blue colors represent the CL site while red/orange colors signify the SL site. Dark colors represent the SG samples. Variation contained in each PC axis are displayed next to each axis.

The total soil C at the SL site increased over the 17-month period in the switchgrass plot (Figure 2a) (r^2^ = 0.12, p < 0.001) and was significantly higher than the fallow (Table 1, p < 0.001, large effect size = 0.4). Switchgrass also had a homogenizing effect for soil C, reducing the overall dissimilarity between samples compared to the fallow plots, which had patchy plant cover. These increases in soil C were occurring evenly across the plots area (Fig. S3a). In contrast, the total soil C content remained constant in the CL switchgrass plot (Figure 2a).

**Figure 2.**
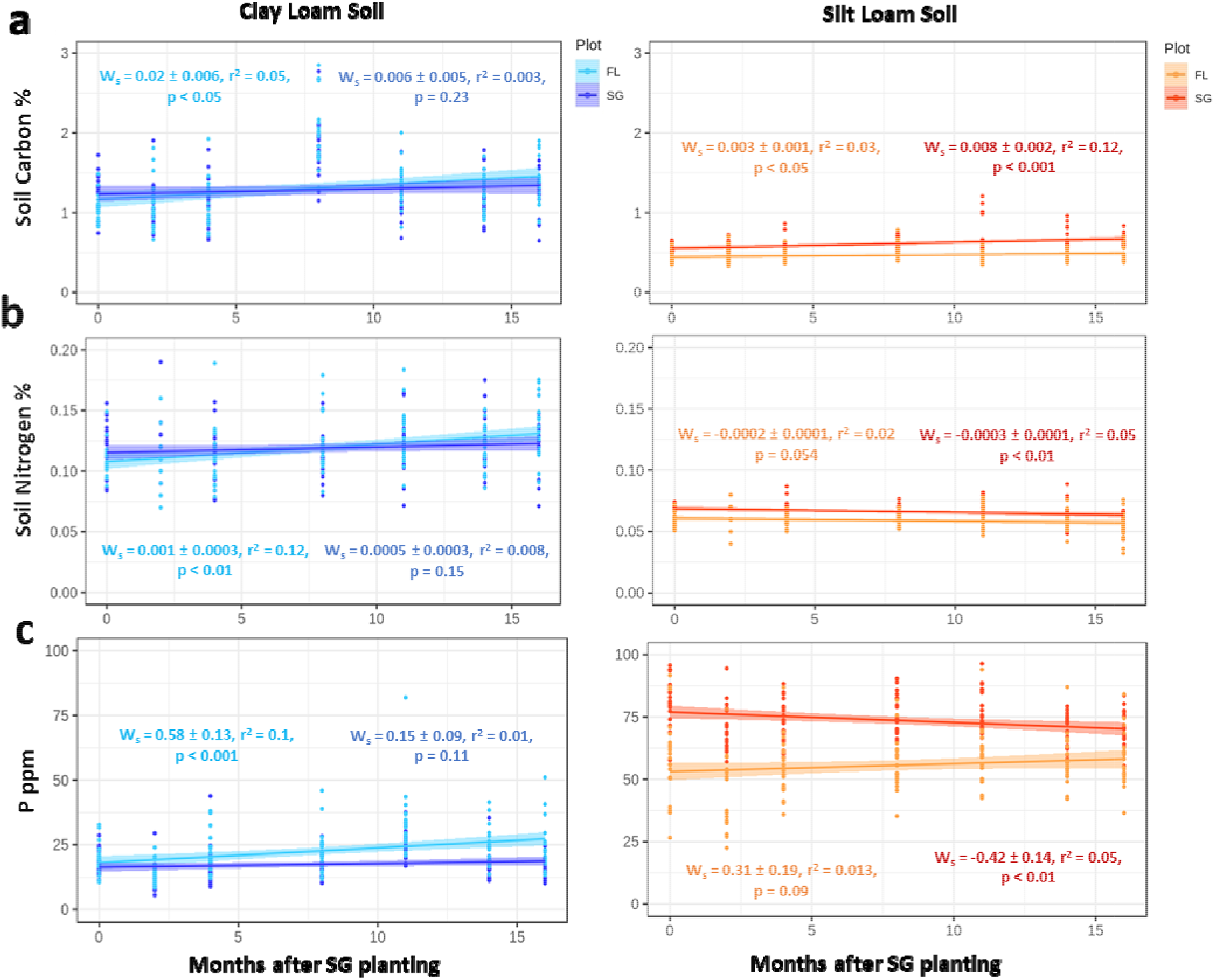
Changes in soil chemistry through two seasons of switchgrass establishment. **a**, Total soil carbon percentages; **b**, Total soil nitrogen percentages; **c**, Concentration of plant available phosphate content in parts per million. The best linear model describing the relationship is presented. W_s_: estimated model slope and associated error. p-values represent the significance of each model. Each time point is comprised of twenty-one replicates per plot.

**Table 1:** Differences in physico-chemical soil properties for each site and treatment after 17-months. Values are mean ± SD values and significance was tested by Kruskal-Wallis rank sum test.

| Variable | Silt Loam<br>Fallow | Silt Loam<br>Switchgrass | Kruskal-Wallis Tests |  |  | Clay Loam<br>Fallow | Clay Loam<br>Switchgrass | Kruskal-Wallis Tests |  |  |
| --- | --- | --- | --- | --- | --- | --- | --- | --- | --- | --- |
|  | Mean ± SD | Mean ± SD | Chi-squared | p | Effect Size | Mean ± SD | Mean ± SD | Chi-squared | p | Effect Size |
| pH | 6.5 ± 0.67 | 6.7 ± 0.95 | 1.32 | 0.25 | - | 5.73 ± 0.44 | 5.85 ± 0.57 | 2.17 | 0.14 | - |
| Soil Moisture (%) | 7.1 ± 4.1 | 8.7 ± 6 | 3.27 | 0.07 | - | 10.4 ± 6.8 | 9.82 ± 5.5 | 0.75 | 0.39 | - |
| Total soil C (%) | 0.47 ± 0.09 | 0.61 ± 0.12 | 118 | < **0.0001\*\*\*** | 0.40 Large | 1.3 ± 0.42 | 1.3 ± 0.36 | 0.86 | 0.40 | - |
| Total soil N (%) | 0.06 ± 0.01 | 0.07 ± 0.01 | 55.4 | < **0.0001\*\*\*** | 0.19 Medium | 0.11 ± 0.02 | 0.11 ± 0.02 | 0.27 | 0.6 | - |
| Phosphorus (ppm) | 55.6 ± 13 | 73.7 ± 9.7 | 128 | < **0.0001 \*\*\*** | 0.44 Large | 22.6 ± 9.7 | 17.5 ± 6.5 | 27.7 | < **0.001\*\*** | 0.095 Medium |
| Nitrate (ppm) | 3.5 ± 3.7 | 2.2 ± 3 | 16.8 | < **0.001\*\*** | 0.06 Small | 8.3 ± 9.8 | 8.9 ± 15 | 2.37 | 0.12 | - |
| Ammonium (ppm) | 13.6 ± 11 | 14.5 ± 10 | 1.73 | 0.19 | - | 18.5 ± 11 | 17.9 ± 11.5 | 0.61 | 0.44 | - |

\*Effect size shown by *epsilon* squared with small (0.01 - < 0.08), medium (0.08 - < 0.26), and large (≥ 0.26) ranges.

Total soil N was significantly higher in the switchgrass plot at SL compared to the fallow plot (Table 1, p <0.0001, medium effect size = 0.19) and these N levels significantly decreased over time (r^2^ = 0.05, p < 0.01) (Figure 2b), coinciding with an increase in the soil N heterogeneity in the plot (r^2^ = 0.12, p < 0.0001) (Fig. S3b). A significant increase in the total soil N was notable in the CL fallow plot (r^2^ = 0.12, p < 0.01) (Figure 2b). Nitrate concentration was significantly reduced for the switchgrass treatment at the SL site (Table 1, p < 0.001, small effect size = 0.06). All sites and plots showed a significant reduction in NO_3_ concentrations over time (Fig. S4) despite an increasing homogeneity (Fig. S3). No significant differences or trends were observed in NH_4_ concentrations during the length of our study at either site (Fig. S3e and S4b).

Total plant available P levels decreased over time in the SL site (r^2^ = 0.05, p < 0.01) (Figure 2c) and the P content homogenized across the plot (Fig. S3c) despite the SL switchgrass treatment having significantly higher total plant available P content compared to the fallow (Table 1, p < 0.0001, large effect size = 0.44). In the CL site, plant available P also decreased in the switchgrass plot compared to the fallow (Table 1, p < 0.001, medium effect size = 0.095, and Figure 2c).

### Differences in estimated root biomass

Field-scale estimates of belowground root biomass (Fig. S2a) showed a large difference in the root biomass between each switchgrass plot and the corresponding fallow (17.8 and 64 times higher for SL and CL, respectively). Root biomass was estimated for each soil layer in kilograms per meter squared and compared to switchgrass estimates. Estimated total root biomass of the switchgrass plots was 16.9 kg/m^2^ for the SL site and 14.1 kg/m^2^ for the CL site, while the fallows were 0.95 kg/m^2^ for SL and 0.22 kg/m^2^ for CL. Generally, root biomass decreased along the soil depth profile for both sites. SL switchgrass site had increased root biomass estimates at lower depths (60-100 cm) than the CL site, which contributed to a slightly higher total root biomass.

### Greenhouse gas (GHG) fluxes at the soil-atmosphere interface

CO_2_ flux at both sites exhibited a similar seasonal trend with the apex of emissions occurring during summer months and the minimum in late Fall/early Winter months. At the SL site, switchgrass treatment led to significantly higher total CO_2_ flux for 29% of the months after switchgrass planting (Wilcoxon p < 0.001, Figure 3a) while the fallow was significantly higher for only 24% of the total months measured. The average CO_2_ flux over the 17-months did not differ in the SL site between switchgrass (6.76 ± 5.23 millimoles·m^2^·hour^−1^) and the fallow (6.87 ± 5.87 millimoles·m^2^·hour^−1^) (Figure 3d). At the CL site, there was a significant difference between treatments (Figure 3d) in the average CO_2_ flux over the 17-month period (p < 0.001) with the switchgrass plot at 9.98 ± 6.04 millimoles·m^2^·hour^−1^ and the fallow at 9.22 ± 6.62 millimoles·m^2^·hour^−1^, although the effect size was small (0.13). When comparing the two sites, CL exhibited significantly higher total soil CO_2_ fluxes for both switchgrass and fallow plots than those measured at SL (Wilcoxon p < 0.001).

**Figure 3.**
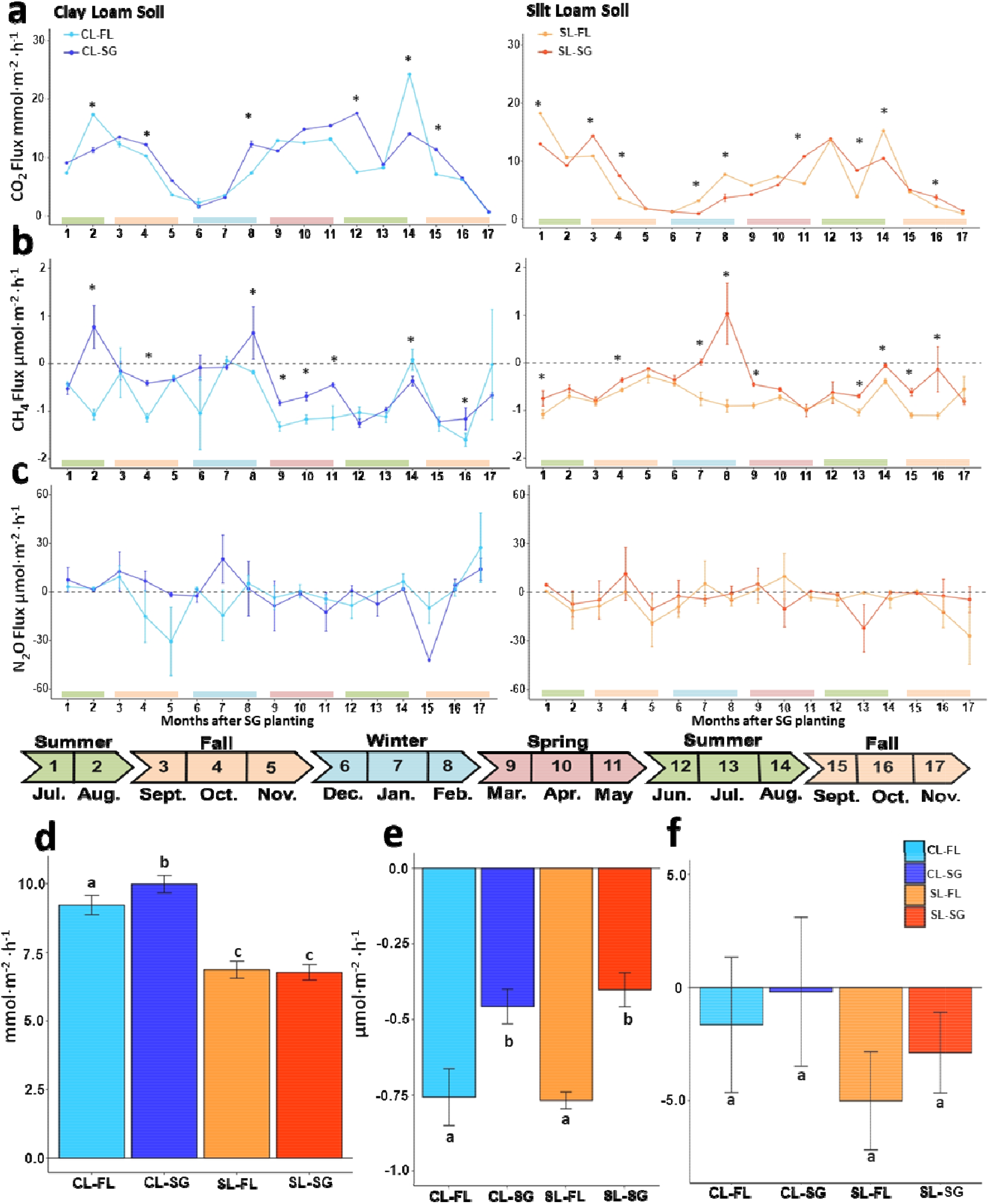
Greenhouse gas (GHG) fluxes during grassland conversion to switchgrass. **a**, **b**, **c**: GHG fluxes at each site over 17 months (mean and standard error estimated using 21 replicates for each time points) for: **a**, carbon dioxide flux; **b**, methane flux; **c**, nitrous oxide. **d**, Average GHG fluxes over 17-months for: **d**, carbon dioxide; **e**, methane flux; **f**, nitrous oxide flux. Different letters and asterisk indicate significant difference between groups by Wilcox sign test with p-value < 0.01.

CH_4_ flux (Figure 3b, Table S3) differed significantly between switchgrass and fallow (Wilcoxon p < 0.001, small effect size = 0.15), with a tendency toward higher CH4 emissions or lower CH_4_ consumption levels in the switchgrass plot observed for 41% of the months (Figure 3b) after switchgrass was planted (41% and 52% for CL and SL, respectively). CH_4_ flux in the fallow was higher only at one time point (14^th^ month after switchgrass establishment). Overall, the 17-month average CH_4_ consumption rate was −0.44 ± 1.07 micromoles·m^2^·hour^−1^ for switchgrass treatments (−0.46 ± 1.08 and −0.41 ± 1.06 micromoles·m^2^·hour^−1^ for CL and SL, respectively) while it reached −0.77 ± 1.15 in the fallow (−0.76 ± 1.78 and −0.77 ± 0.53 and micromoles·m^2^·hour^−1^ for CL and SL, respectively) (Figure 3e, Table S3). Together, a significant (p < 0.05, a small effect size = 0.14) switchgrass treatment effect on reducing CH_4_ consumption rates was observed at both sites. No significant differences were found for N_2_O flux between the switchgrass (−0.26 ± 2.55 micromolesom·^2^·hour^−1^ at CL and −2.88 ± 2.09 micromolesom·^2^·hour^−1^ at SL) and fallow plots (−1.65 ± 2.5 micromoles per m^2^ per hour at CL and −5.01 ± 2.16 micromoles per m^2^ per hour at SL) at either site (Figure 3c, Table S3) over the 17-months of observations (Figure 3f).

### Microbial community dynamics

Microbial alpha diversity, calculated as the OTU richness, responded in a site-specific manner to switchgrass cultivation. In the SL site, OTU richness was significantly higher in the switchgrass plot (Table S3, p < 0.0001, Medium effect size = 0.38). OTU richness did not change over time in the SL switchgrass plot (Figure 4a) but increased in the fallow plot (p < 0.001), despite a decrease in phylogenetic diversity (PD) (p < 0.05, Figure 4a,b). At the CL site, species richness decreased significantly over time in both switchgrass (p < 0.01) and fallow plots (p < 0.001). For PD, this decay was observed only in the switchgrass plot (p < 0.01). Chao1 and Shannon index showed similar trends per site over time (Fig. S5).

**Figure 4.**
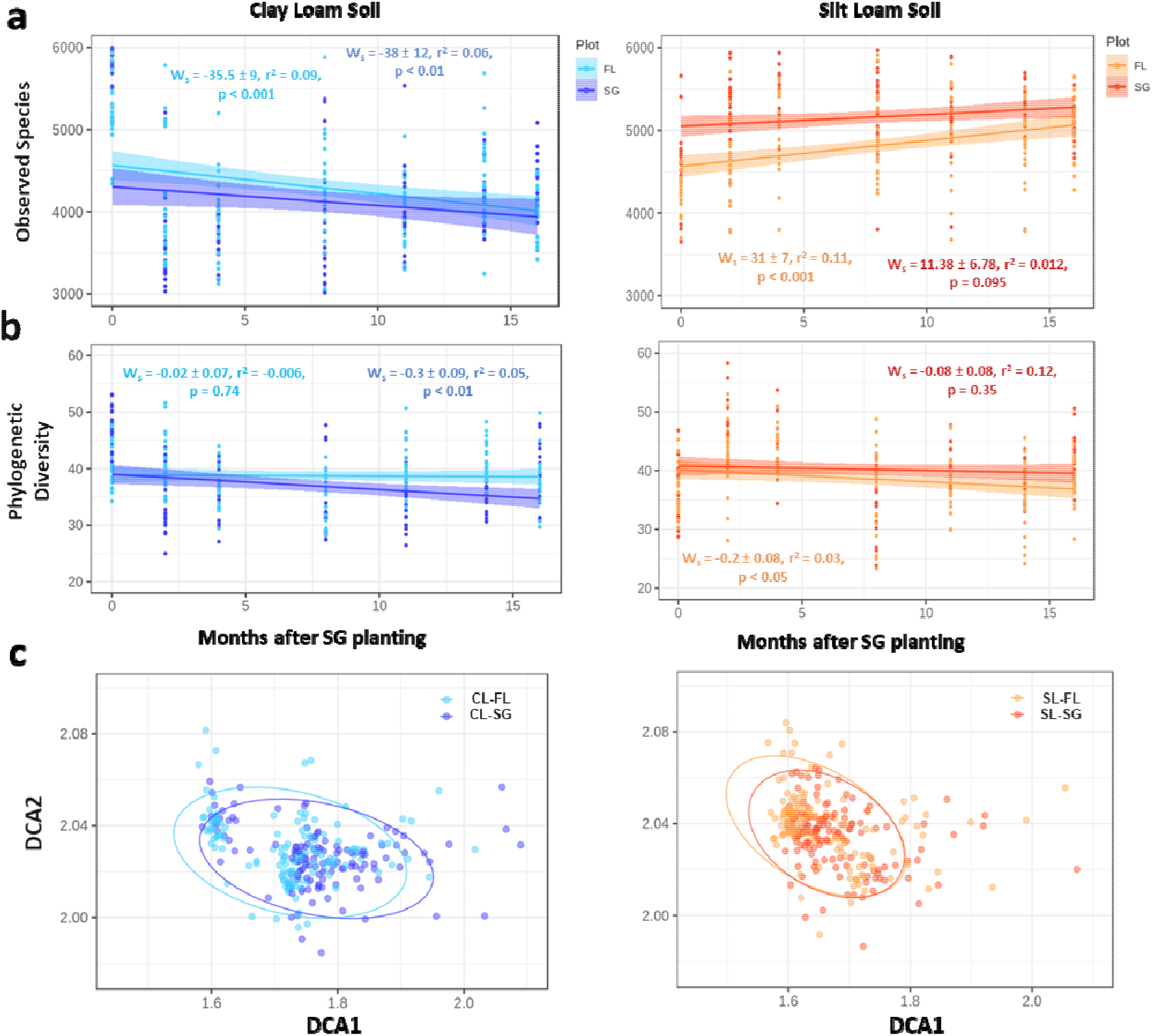
Changes in microbial diversity and structure in response to switchgrass planting. **a**, Number of observed species through time; **b**, Phylogenetic diversity. **c**, Detrended correspondence analysis of the 16S community separated by site for all time points and plots. Significant differences were found between sites, plant cover types, and through time (PERMANOVA, p < 0.01). Dark colors represent the switchgrass samples.

We observed significant differences in the bacterial community structure (beta diversity) between sites, plant cover type, and over time (Figure 4c, PERMANOVA, p < 0.01, Table S4). Relative abundance of major phyla showed large changes from the initial planting/before soil tillage and two months after the experiment began (Figure 5). At all sites, at least five abundant phyla exhibited changes in relative abundance. Firmicutes phyla relative abundance (0.6 – 0.14 %) changed over the course of the experiment in both fallow plots. The structure of microbial communities from the switchgrass plots appeared less variable than their corresponding fallows. In the CL site, the strongest differences in dominant phyla relative abundance between plots (switchgrass vs fallow) were observed at eight and fourteen months after switchgrass planting (Feb. 2017 and Aug. 2017, Table S5). After eight months, seven phyla (Actinobacteria, Bacteroidetes, Chloroflexi, Deinococcus-Thermus, Firmicutes, Plactomycetes, and Verrucomicrobia) exhibited different abundance between treatment, while only four phyla (Actinobacteria, Chloroflexi, Cyanobacteria, and Deinococcus-Thermus) were different after fourteen months. For the SL site, the largest shifts in community composition occurred in the last two time points, *i.e*. fourteen and sixteen months after switchgrass establishment. After fourteen months, three phyla were significantly different between treatment (Acidobacteria, Bacteroidetes and Deinococcus-Thermus) and after sixteen months four groups were significantly different (Acidobacteria, Bacteroidetes, Cyanobacteria, and Verrucomicrobia).

**Figure 5.**
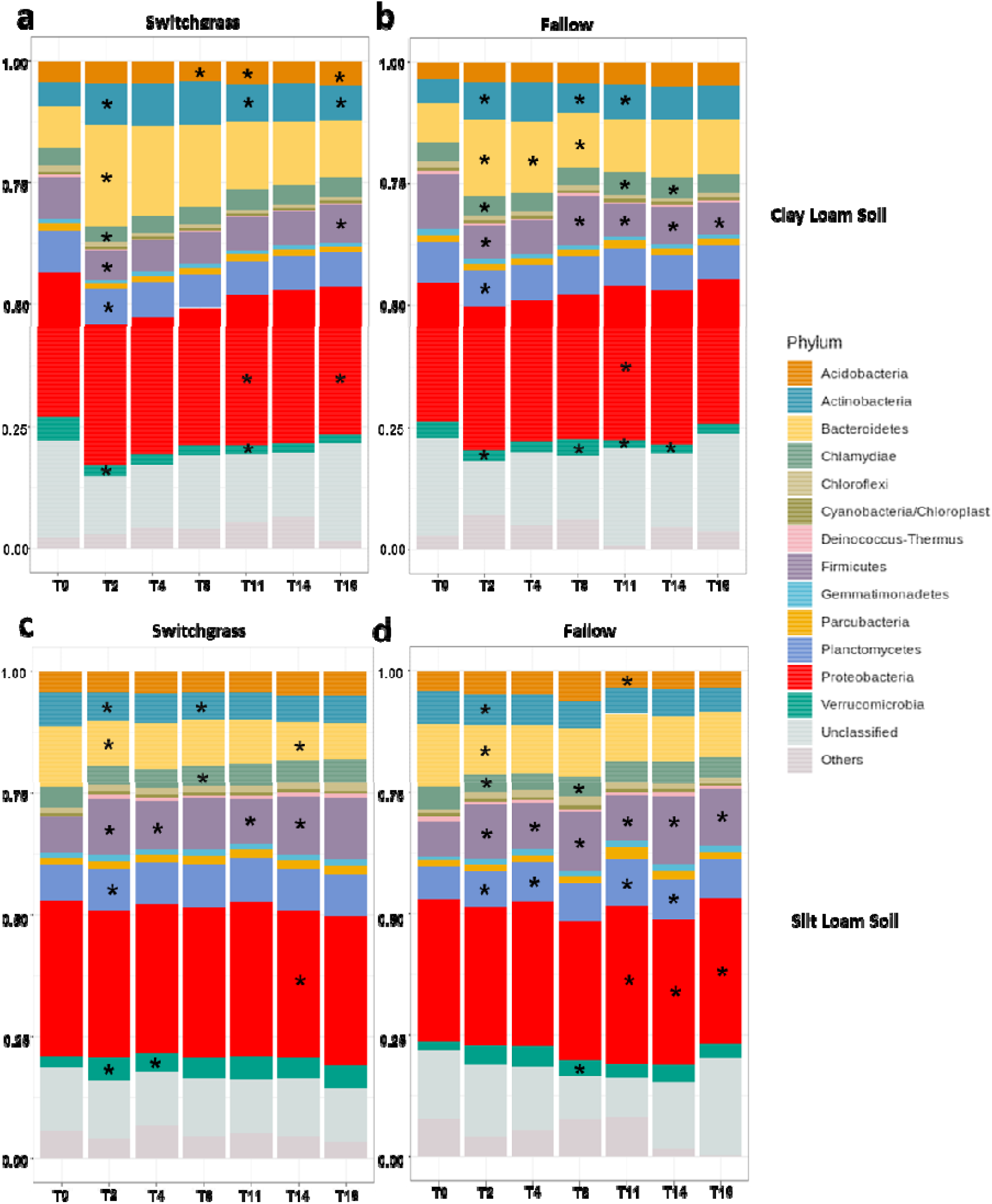
Changes of relative abundance for major phyla. Taxonomic identity was determined with the RDP classifier at 80% sequence match criteria. OTU table was trimmed by abundant OTUs (> 0.001%). Difference between time points within each plot for: **a**, Clay-loam switchgrass (CL-SG) plot; **b**, Clay-loam fallow (CL-FL) plot; Silt-loam switchgrass (SL-SG) plot; Silt-loam fallow (SL-FL) plot. Significant differences between the pervious time point for each group denoted by asterisk (*) symbols within each phyla bar.

Canonical correspondence analysis (CCA) was used to link environmental variables to the microbial community (Figure 6). A clear separation between microbial communities from the two sites was observed. Microbial communities from the SL site were correlated with plant available P and soil pH, while CL communities were associated with total soil N, NH_4_, and NO3. In addition, fallow communities from CL were dispersed, while switchgrass communities at this site were clustered by N source or along a soil moisture profile.

**Figure 6.**
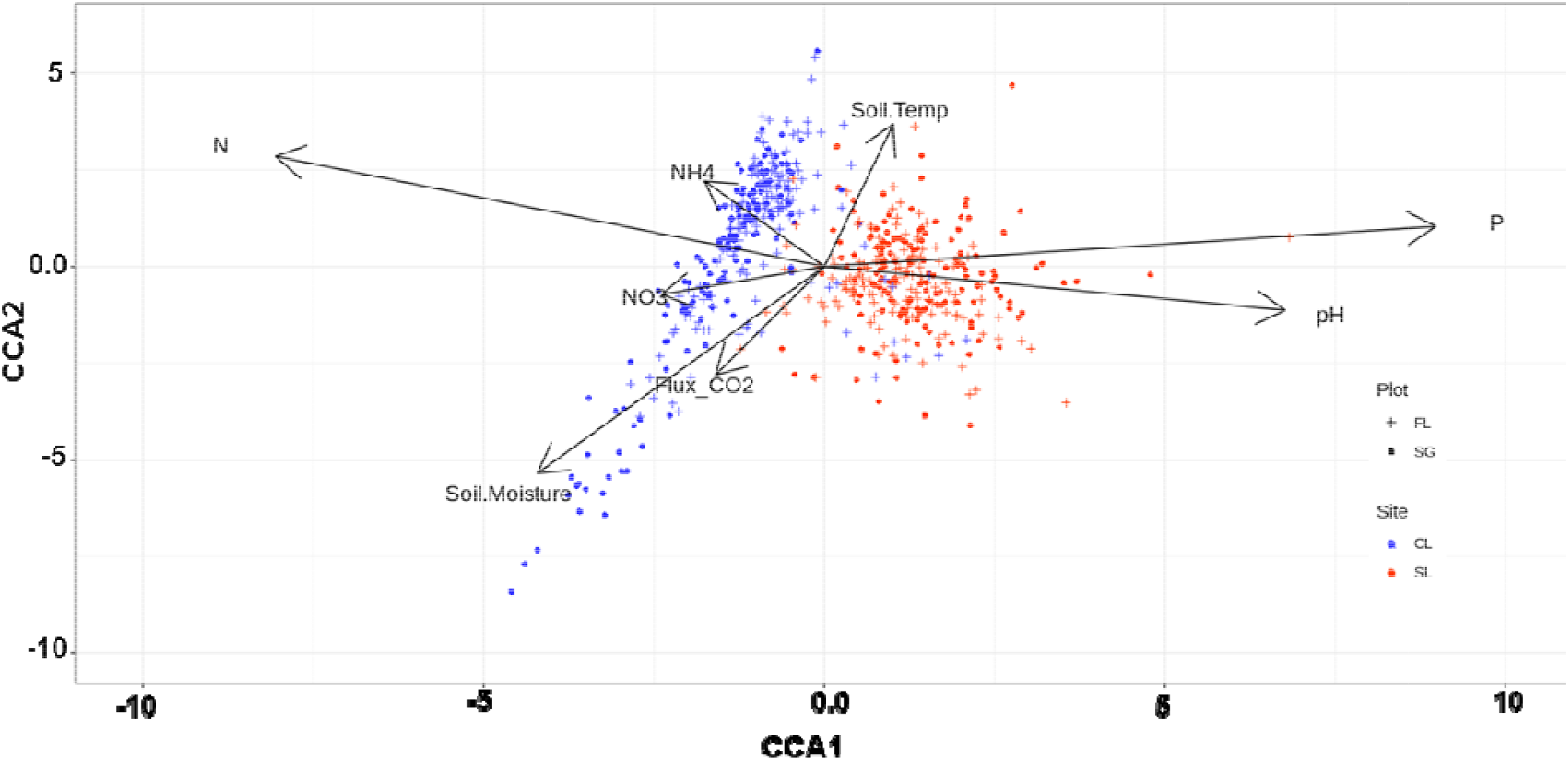
Relationships between environmental factors and microbial communities structure. Canonical correspondence analysis (CCA) linking microbial communities structure with environmental variables. Samples are shown by plot and site type with significant environmental variables shown in black arrows.

### Structural equation model

Structural equation modeling (SEM) was used for an in-depth analysis of the direct and indirect effects of the environmental drivers on CO_2_ and CH_4_ flux for both sites. Correlations between all variables are shown in the correlogram Figure S6. For CO_2_ fluxes (Figure 7a) the model confirmed the importance of the site effect on soil geochemistry and microbial communities, with the strongest direct effects (based on standardized coefficient) being directed from the site towards total C (β = −0.95, p < 0.001), total N (β = −0.94, p < 0.001), P levels (β = 0.88, p < 0.001), and microbial alpha diversity (β = 0.52, p < 0.001). Plant available P tended to influence the levels of C and N in the system, and these three components of soil have significant effects on CO_2_ fluxes. Other important variables influencing CO_2_ fluxes included soil temperature (β = 0.69, p < 0.001), moisture (β = 0.27, p < 0.001) and microbial biomass (β = - 0.18, p < 0.001). This later variable appeared mostly dependent on N content (β = 0.56, p < 0.001).

In the CH_4_ model (Figure 7b), which was focused on the switchgrass plots, the site effect appeared less pronounced and mostly directed toward P levels (β = −0.29, p < 0.001) and microbial alpha diversity (β = 0.62, p < 0.001). Although soil temperature did not have a very strong influence on CH_4_ fluxes (β = −0.19, p < 0.001), it was still important in this model through many direct effects on P (β = 0.07, p < 0.001), NO_3_ (β = 0.23, p < 0.001), pH (β = −0.22, p < 0.001), microbial diversity (β = 0.11, p < 0.001), and moisture (β = −0.39, p < 0.001). Overall, we were able to show that CH_4_ fluxes depended on a combination of soil chemical properties (P and NO_3_) along with physical (temperature and moisture) and biological (microbial diversity) characteristics.

**Figure 7.**
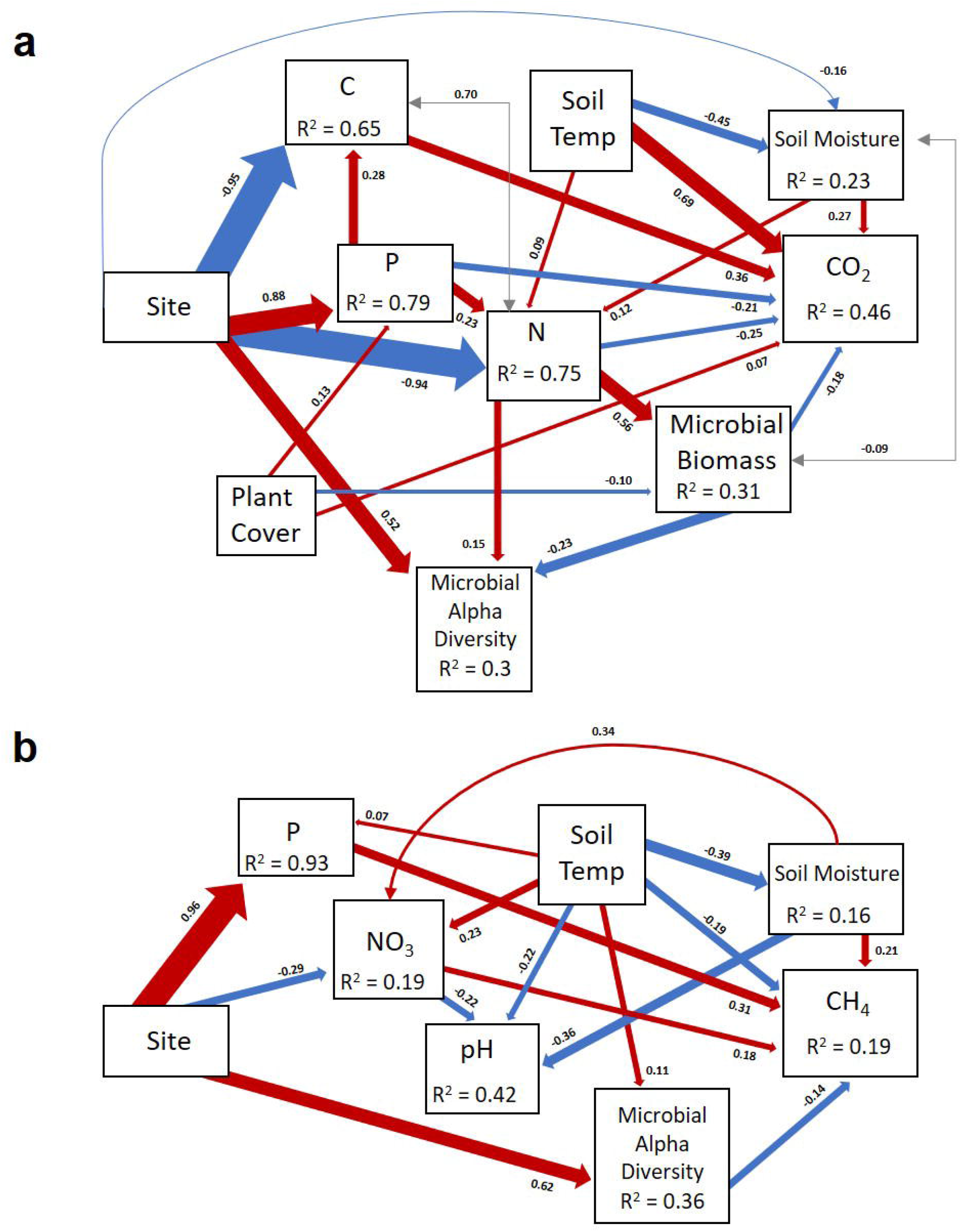
Structural equation modeling showing the relationships among environmental variables and GHG fluxes. **a**: Model for total carbon dioxide flux generated from the seasonal data (*χ*^2^ = 25.806, d.f. = 18, *P* = 0.104, *n* = 588). **b**: Model for methane flux generated from seasonal data of switchgrass plots only (*χ*^2^ = 10.116, d.f. = 9, *P* = 0.341, *n* = 294). Red and blue arrows represent significant (p < 0.05) positive and negative pathways, respectively. Numbers near the pathway arrows indicate the standard path coefficients (β). Width of the arrows are proportional to the strength of the relationship. Gray arrows represent residual correlations accounted for in the model. Plant Cover = Switchgrass (positive) or mixed annual grassland plant cover (negative) at the plot; Site = SL (positive) or CL (negative) soil site; CO_2_ = total soil carbon dioxide flux; Soil Temp = soil temperature at a depth of 10 cm for bare soil in degrees Celsius; Soil Moisture = gravimetric per cent soil moisture; P = plant available phosphorus content; Microbial Alpha Diversity = number of observed bacterial species per sample; NO_3_ = nitrate concentrations; CH_4_ = methane flux; and pH = soil pH.

## Discussion

Our study illustrates that soil type is a key component of switchgrass’ ability to improve soil quality and appears crucial for its use as a sustainable bioenergy feedstock. It was hypothesized that switchgrass would increase (i) topsoil C levels and (ii) increased CO_2_ respiration -with (iii) little effect on CH_4_ emissions and N_2_O flux, but also (iv) decreased the bulk soil microbial species richness relative to annual mixed grassland communities. While C accrual, CO_2_ respiration, and microbial richness reductions were site-dependent in our study, switchgrass significantly altered the soil CH_4_ sink capacity at both sites by reducing the overall CH_4_ consumption rates. Although the CH_4_ emission rates reported here were expectedly lower than those reported for natural systems with anaerobic, water-logged conditions like peatlands (Dise, 1993) and wetlands (Bridgham et al., 2013; Bartlett and Harriss, 1993), our findings challenged the notion that CH_4_ emissions has negligible effects on GHG budgets during marginal land transitions to switchgrass row-cropping (Monti et al., 2012). However, comprehensive GHG budgets along with spatial-explicit modeling of soil and plant C stocks should be considered to fully evaluate the net effect of land type conversion at these prairie sites.

### Soil type dictates the effects that switchgrass has on geochemistry

Our study revealed significant site-level differences to switchgrass establishment on soil C accrual, total soil N levels and depletion of soil P content. The CL site with higher relative nutrient content showed little change over time (17 months) for any of the soil geochemical parameters. Ma et al., 2000 reported changes in soil geochemistry by switchgrass cultivation in clay-loam soils were only detected after longer periods of time (over a decade). Therefore, prolonged sampling at our CL site will improve assessments of switchgrass-induced soil C changes. Nitrate contents (Figure S4a) significantly declined with time, which may suggest assimilation by the switchgrass or an increased activity of denitrifying bacteria. It is also notable that the crop was grown under natural conditions without applying any chemical fertilizers.

The SL site showed significant increases in soil C content for the top-soil layer (27% higher total C after two growing seasons) over the course of switchgrass establishment. This is consistent with estimates of switchgrass systems repaying their C debt in a relatively short amount of time (Abrah et al., 2019). However, switchgrass led to significant depletions in soil N and P contents at the SL site over time, though these values were higher than the fallow from the beginning of our experiment. One explanation for this observation is the increased below ground root biomass estimates being larger for the SL site at lower depths. Greater investment of belowground root biomass may reflect the higher plant available P conditions at this site, allowing switchgrass to extend deeper into the subsurface soil for water or micronutrient availability.

Seasonal sampling of NH_4_-N sources was not sufficient in explaining large seasonal variations observed over the time course of our experiment. For example, a spike in soil NH_4_ levels (Figure S4b) was detected during October of 2016 and June of 2017, which could be the signature of episodic N fixation events occurring in switchgrass during/before flowering as reported previously (Roley et al., 2019). However, our resolution in sampling this geochemical parameter is not sizeable enough to adequately explain these anomalies.

### Microbial community shifts under switchgrass establishment

Microbial community diversity and composition at each site had differential responses to switchgrass establishment. Broadly, alpha diversity measures in CL decreased over time and revealed a higher amount of clustering and similarity in the overall community structure compared with the fallow. Analogous to secondary plant successional dynamics, the microbial community at the CL site may be more influenced by the change from short rooted annuals to the monoculture of deep-rooted perennial switchgrass, causing a loss in microbial diversity (Cline and Zak, 2015). For SL, switchgrass cultivation significantly increased the Shannon index over time and caused the community composition to shift away from the fallow. This may be indicative of the improvements of soil quality, which changed the functionality of the microbial community due to the influence of switchgrass on increases in soil C levels (Leff et al., 2015).

Microbial community structure was altered by switchgrass establishment (Table S4) and through time at each of the sites relative to the fallow soil communities. These changes in community structure may reflect different survival strategies that switchgrass may employ in the recruitment of specific taxa to its rhizosphere based of differences between the geochemistry of the two sites. Investigations into rhizosphere microbiome succession during establishment may provide insights into direct plant-microbe interactions that facilitate switchgrass establishment in these nutrient-poor soils.

### Factors controlling soil GHG flux

Contrary to our hypothesis, CO_2_ respiration was significantly enhanced by switchgrass establishment only at the CL site. We expected higher root respiration and the potential for deep C mineralization to enhance soil respiration at both sites after switchgrass establishment (Shahzad et al., 2018; Fontaine et al., 2007). The CL site had an overall higher total CO_2_ emission rate during our field monitoring. This response may be mediated by the relatively higher preexisting C nutrient conditions found at this clay-loam site (Kang et al., 2016) as root biomass levels were estimated to be similar at each site. This illustrates a benefit in site selection by soil type in minimizing CO_2_ released during land conversion.

Nitrous oxide fluxes did not show any significant effect between either site or plant cover type. Thus, we did not see an effect of switchgrass or site on nitrous oxide fluxes. However, nitrous oxide fluxes were marked by high variability, both seasonally and spatially within each plot. Efforts were made to correlate rainfall events to reduce noise in the flux signal, but a substantial limitation was our six-minute window of continuous measurements per sampling event. A longer period of trace gas sampling may have resulted in a more stable signal with less variability for nitrous oxide fluxes.

Methane flux monitoring showed a significant reduction in CH_4_ consumptions at both sites with switchgrass introduction and cultivation. Although CH_4_ emission rates were low and measured at only a few time points, consistently lower CH_4_ consumption rates were observed throughout the monitoring period of our experiment. Total methane consumption rates for switchgrass plots were reduced by 47% and 39% compared to corresponding fallow sites for CL and SL, respectively. This could reflect considerable differences in the net C budget and fluxes for these switchgrass sites. In the future, GeoChip-based functional microarray (Shi et al., 2019) as well as, RT-qPCR of CH_4_ monooxygenase and methyl-coenzyme M reductase genes may help provide us with specific linkages between microbial functionality potential and our reported CH_4_ emissions at key time points during our experimental monitoring.

## Conclusion

Overall, soil C levels increased by 27% during the 17 months experiment in the site with the lowest nutrient content (silt loam, SL) while they remained consistent in the clay loam (CL) site. Switchgrass significantly affected total CO_2_ respiration at the CL site, but not at the SL site compared to the annual mixed grassland community fallows and showed a difference in the site level emissions. Grassland conversion to switchgrass reduced the annual CH_4_ consumption by 39 to 47%, implying that methane fluxes should be accounted for in C budgets to reach a sustainable cultivation of switchgrass. Switchgrass establishment had a significant influence on the microbial community composition over time. Our SEM analysis indicated that soil temperature and moisture were strong environmental drivers of the soil the CO_2_ and CH_4_ flux at each site. Considerations on soil type and nutrient conditions should be made for the selection of future sites suitable for large-scale bioenergy cultivation of that meets objectives for terrestrial C sequestration and improved soil fertility.

## Acknowledgements

We would like to thank the Oklahoma Mesonet environmental monitoring network, and particularly Bradley Ilston, for the use of their weather monitoring stations for this work. We deeply appreciate all the help from the auxiliary staff and field hands from the Nobel Research Institute, who aided in us in collecting data and maintaining the field sites for this project. A. Escalas and C. Bates would like to give a special thanks to Randy Freeman, who helped a lot for making sample collection and processing possible. We would also like to thank the following for their contributions in field sampling and molecular work: Zhigang Wang, Yuguang Zhang, Ning Hu, Yan He, Zhongfang Li, Qian Li, Jinyu Hou, Xiubin Ke, Juan Ling, Zheng Gao, and Daniel Curtis. C. Bates would also like to thank Cheryl Bates for edits and Tanya Ball for support during this project. This research is supported by the U.S. Department of Energy Office of Science, Office of Biological and Environmental Research Genomic Science program under the award number DE-SC0014079 to the UC Berkeley, Nobel Research Institute, University of Oklahoma, the Lawrence Livermore National Laboratory, and the Lawrence Berkeley National Laboratory.

---

**Fig. S1.**
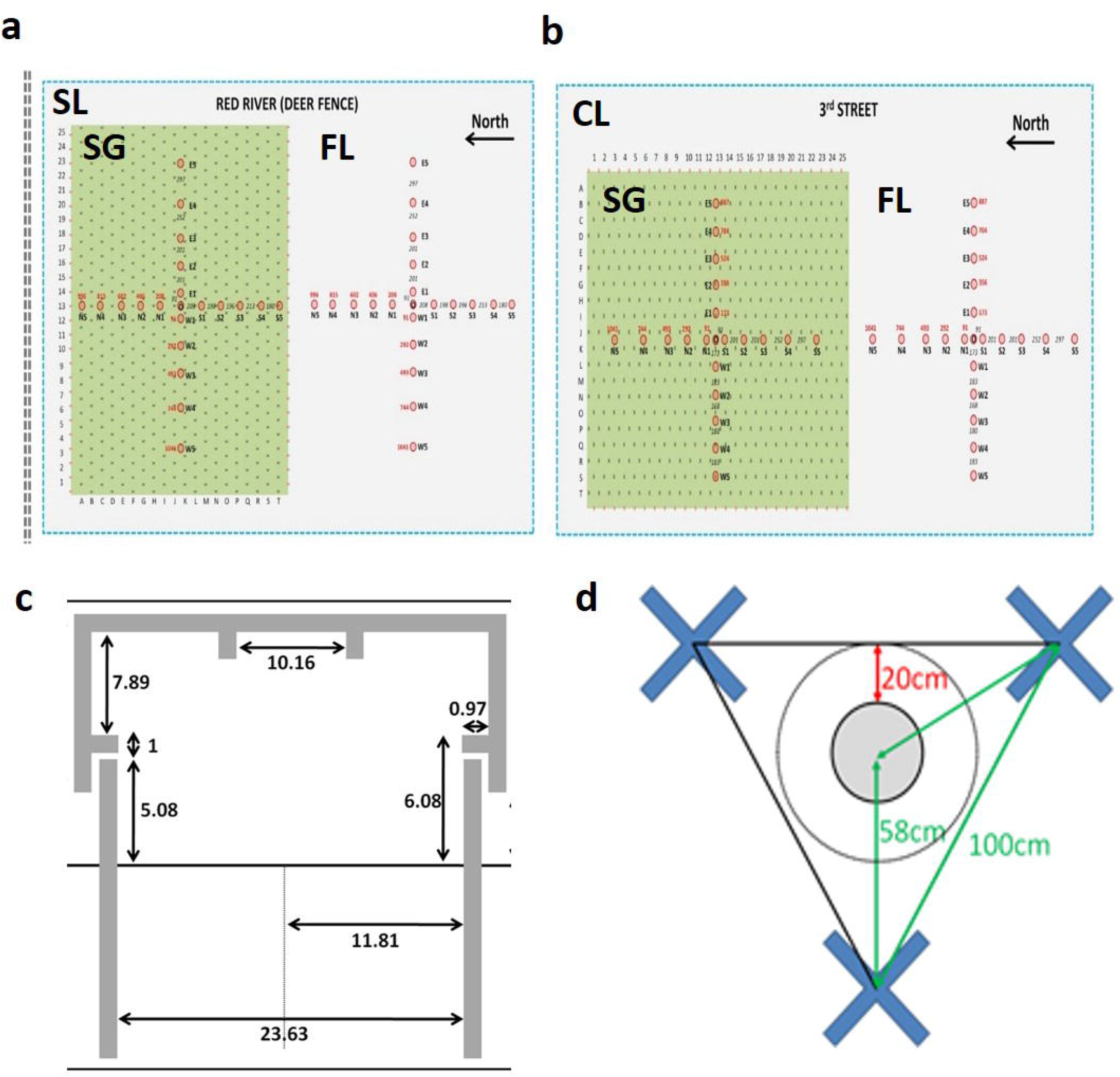
Layout for the field sites, collar positions, and soil sampling details: **a,** layout of the SL field site and location of trace gas collars for each plot; **b**, layout of the CL field site and location of the trace gas collars for each plot; **c,** diagram of trace gas collar and chamber dimensions; **d,** diagram of collar position in relation to switchgrass plants and soil sampling details. Blue X’s represent switchgrass plants and the gray circle represents the trace gas collar.

**Fig. S2.**
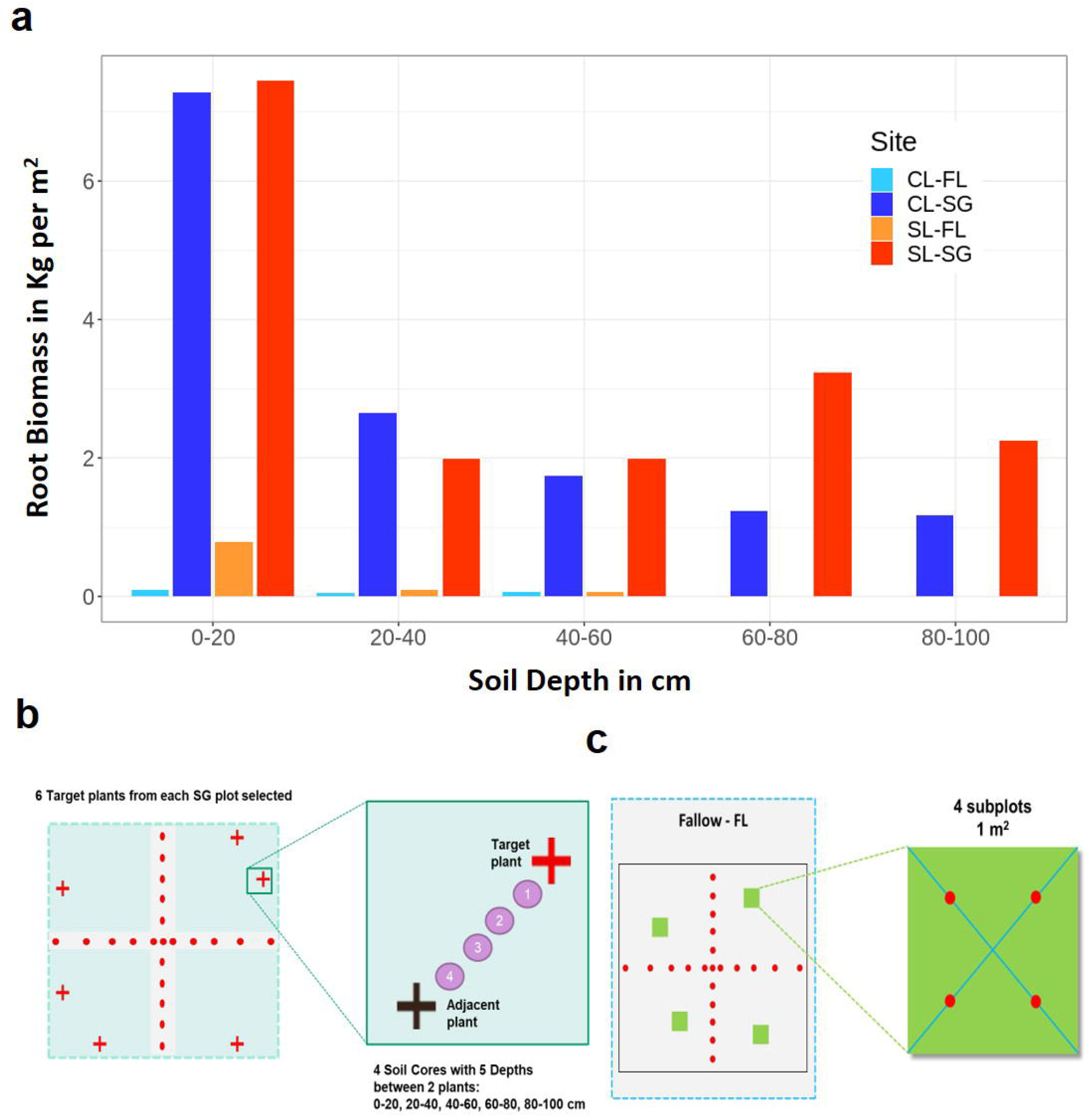
Root biomass estimates and methods for: **a**, difference between fallow and switchgrass plots for estimated root biomass by depths; **b**, graphic for how switchgrass plants were selected for root biomass estimation; **c**, graphic for how fallow root biomass estimation was conducted using four 1 m^2^ subplots in each quadrant of the plot.

**Fig. S3.**
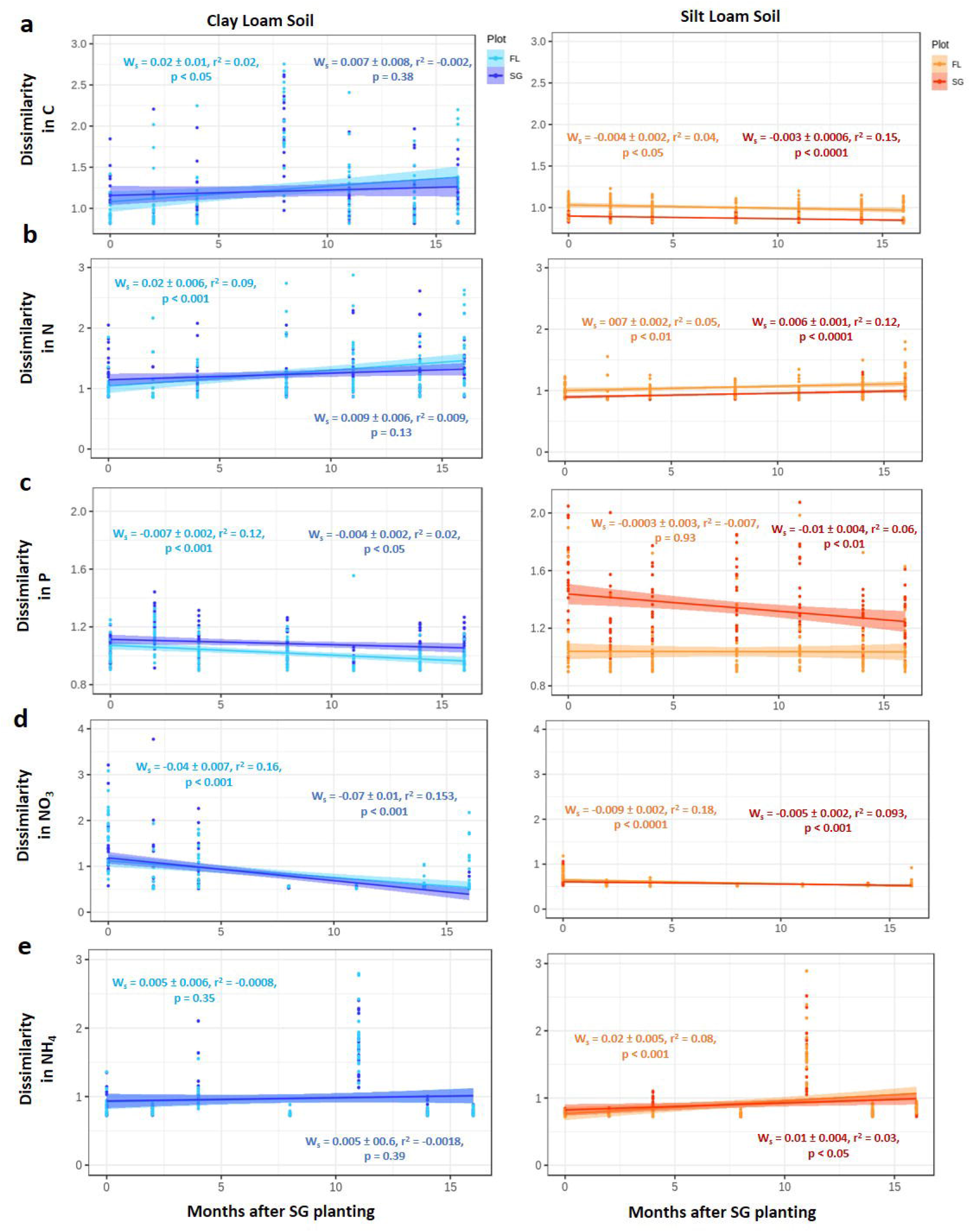
Dissimilarity between soil chemistry samples through the seasons at each site between SG and FL plots for: **a**, dissimilarity in soil carbon samples; **b**, dissimilarity in soil nitrogen content; **c**, dissimilarity in plant available phosphate content; **d**, dissimilarity in nitrate content; **e**, dissimilarity in ammonium. Ws is the slope of each line and the error associated with each slope while p-values represent the significance of each trend line. Each time point is comprised of twenty-one replicates per plot.

**Fig. S4.**
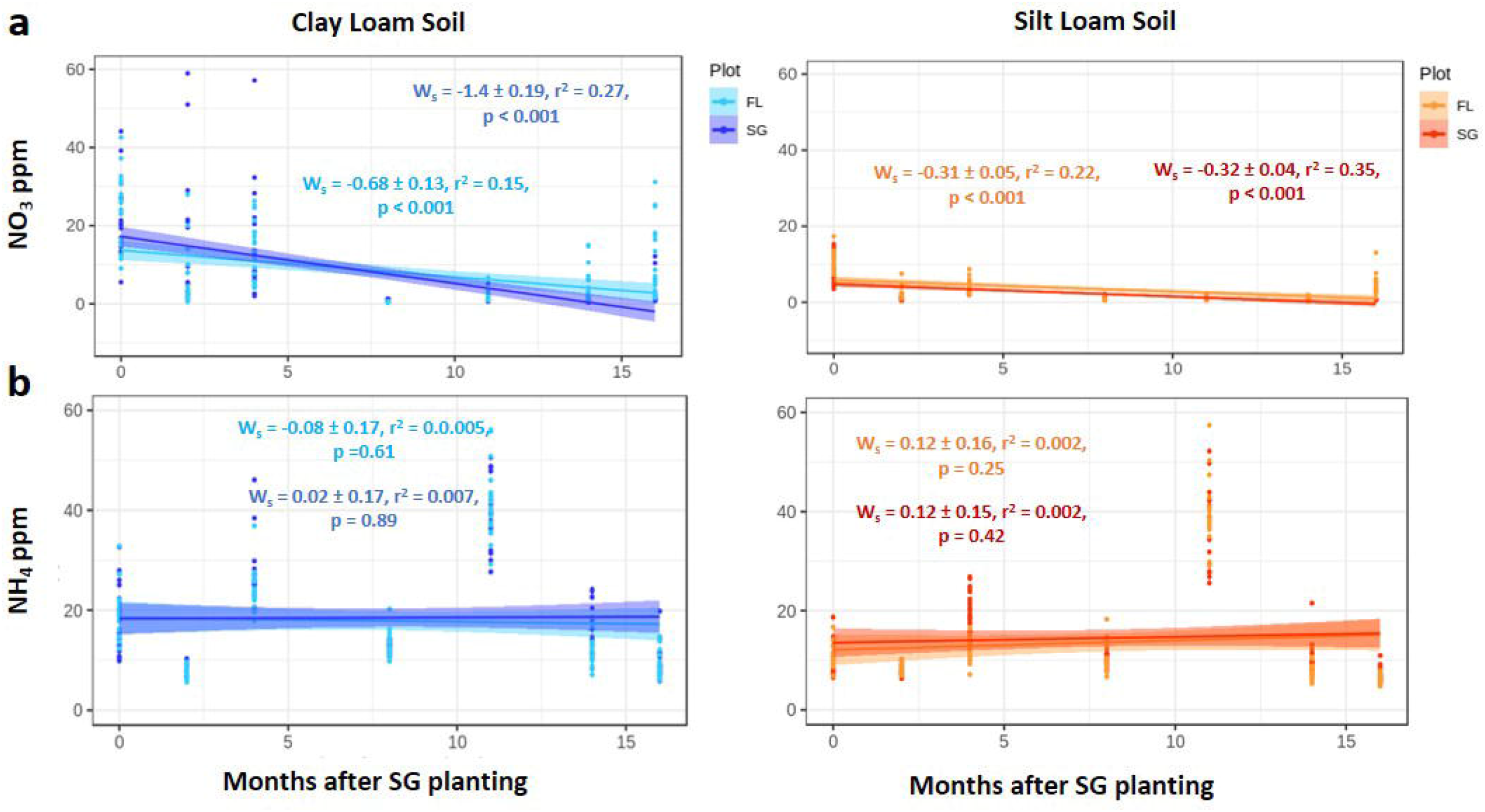
Changes in soil chemistry through the seasons at each site between SG and FL plots for: **a**, concentration of nitrate content; **b**, ammonium content. Ws is the slope of each line and the error associated with each slope while p-values represent the significance of each trend line. Each time point is comprised of twenty-one replicates per plot.

**Fig. S5.**
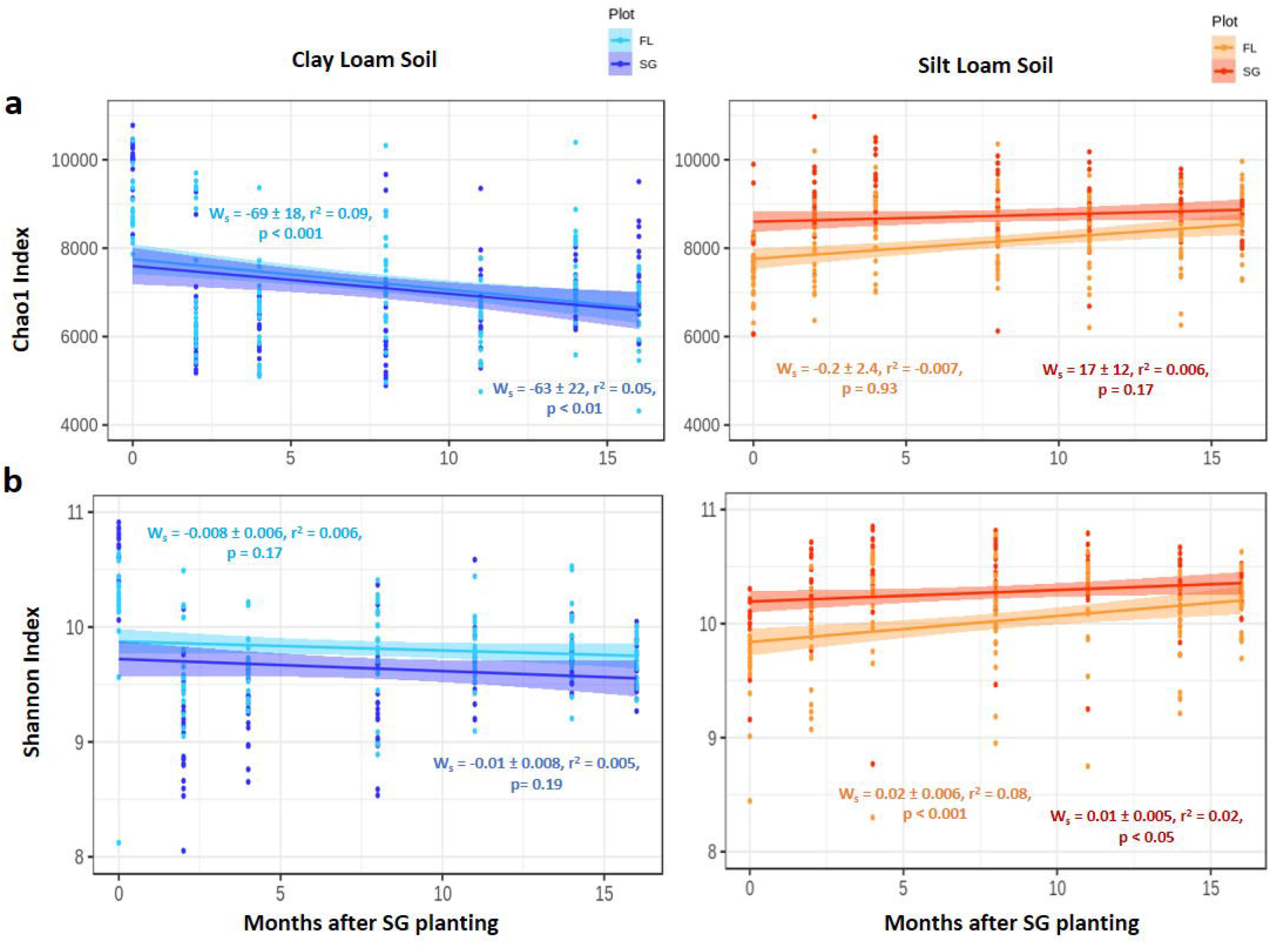
Change in microbial alpha diversity measures through the seasons at each site between SG and FL plots for: **a**, chaol index; **b**, Shannon index. Ws is the slope of each line and the error associated with each slope while p-values represent the significance of each trend line. Each time point is comprised of twenty-one replicates per plot.

**Fig. S6.**
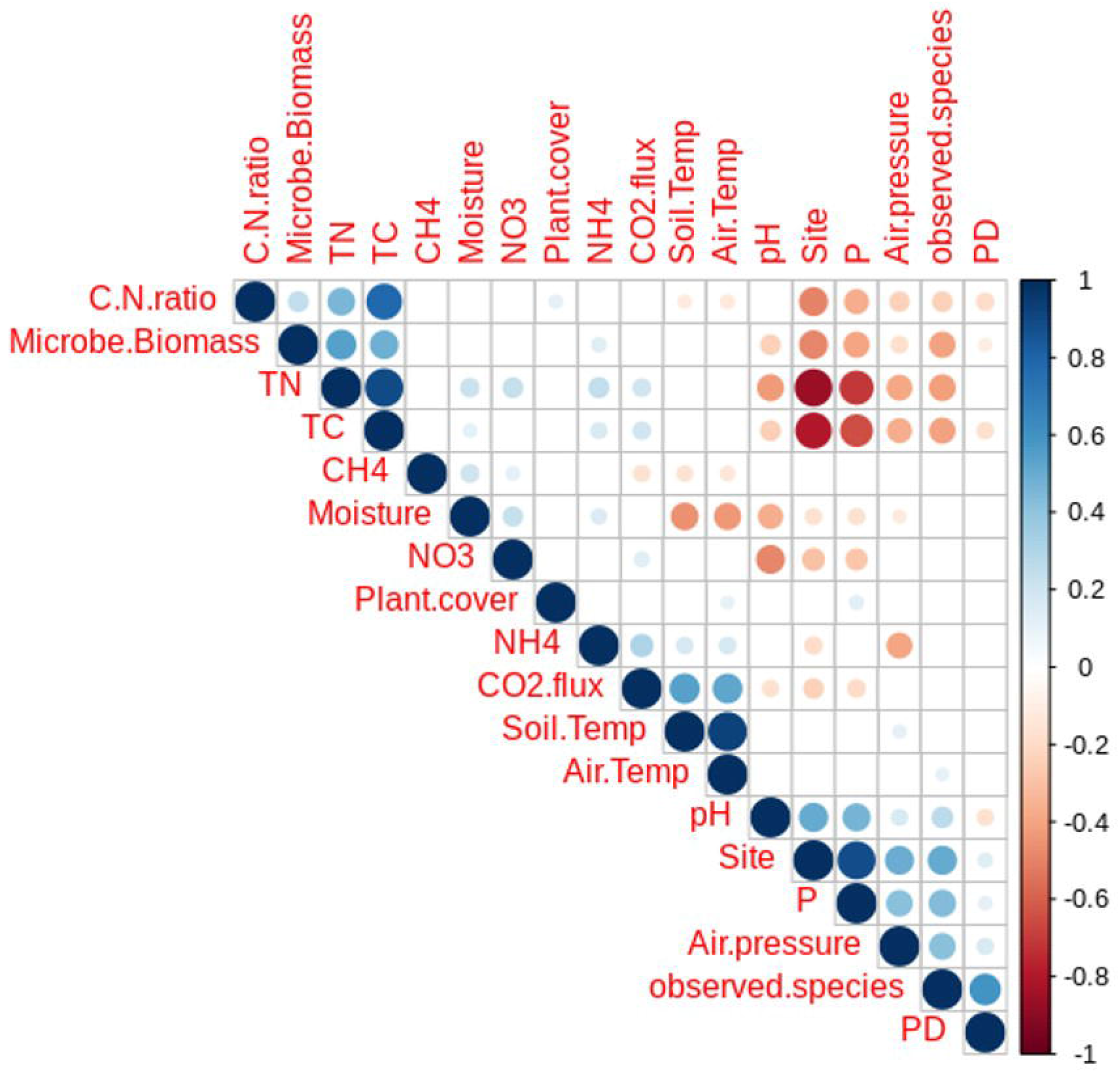
Correlation plot between environmental, microbial, and soil geochemistry variables. Blue circles indicate significant positive correlations while red circles indicate negative correlations. Larger darker circles indicate a more significant and stronger correlation between variables. Positive correlations for site represent SL site while negative correspond to the CL site. ‘Microbe.Biomass’ was estimated using DNA concentrations after soil extractions, ‘observed.species’ is the total species richness (alpha diversity) of the samples. ‘PD’ represents the whole tree phylogenetic diversity of the plots. Positive correlations for ‘Plant.cover’ represent switchgrass plots and negative for fallow plots.

