## Supplementary material for "Conversion of marginal land into switchgrass conditionally accrues soil carbon and reduces methane consumption": Supp. Figures Document

Bates et al. Supplementary Figures and Tables

**Table Captions**

**Table S1.** Seasonal data set for environmental factors, soil chemistry, and GHG fluxes by sample ID.

**Table S2.** Identification of variable names in found in Table S1.

**Table S3.** GHG flux and community metrics differences by Wilcox Sign Ranked Test and Mann-Whitney U test for effect size between switchgrass (SG) and fallow (FL) plots.

**Table S4.** PERMANANOVA for 16S beta community structure by different treatment effects based off 999 permutations.

**Table S5.** p-values from t-test between relative abundances of major phyla for SG vs FL groups of each soil type by time point. p values adjusted by conservative Bonferroni correction method.

| **Table S1.** Seasonal data set for environmental factors, soil chemistry, and GHG fluxes by sampleID. | | | | | | | | | | | | | | | |
| --- | --- | --- | --- | --- | --- | --- | --- | --- | --- | --- | --- | --- | --- | --- | --- |
| SampleID | SampleName | Date | Year | Season | Month | Months_After_SG | Site | Plot | CollarPosition | Direction | Ring | Subplot | pH | Moisture_Grav | RELH |
| 43 | CL_T0_SG_E1 | 20160718 | 2016 | Summer | July | 0 | CL | SG | E1 | E | 1 | CL-SG | 5.57 | 12.08053691 | 67 |
| 44 | CL_T0_SG_E2 | 20160718 | 2016 | Summer | July | 0 | CL | SG | E2 | E | 2 | CL-SG | 5.59 | 7.643312102 | 67 |
| 45 | CL_T0_SG_E3 | 20160718 | 2016 | Summer | July | 0 | CL | SG | E3 | E | 3 | CL-SG | 5.52 | 9.347826087 | 68.25 |
| 46 | CL_T0_SG_E4 | 20160718 | 2016 | Summer | July | 0 | CL | SG | E4 | E | 4 | CL-SG | 5.42 | 11.52993348 | 68.875 |
| 47 | CL_T0_SG_E5 | 20160718 | 2016 | Summer | July | 0 | CL | SG | E5 | E | 5 | CL-SG | 5.42 | 9.649122807 | 70.125 |
| 48 | CL_T0_SG_N1 | 20160718 | 2016 | Summer | July | 0 | CL | SG | N1 | N | 1 | CL-SG | 4.88 | 8.893709328 | 64 |
| 49 | CL_T0_SG_N2 | 20160718 | 2016 | Summer | July | 0 | CL | SG | N2 | N | 2 | CL-SG | 5.47 | 8.189655172 | 51 |
| 50 | CL_T0_SG_N3 | 20160718 | 2016 | Summer | July | 0 | CL | SG | N3 | N | 3 | CL-SG | 5.28 | 4.31211499 | 48 |
| 51 | CL_T0_SG_N4 | 20160718 | 2016 | Summer | July | 0 | CL | SG | N4 | N | 4 | CL-SG | 5.17 | 7.188160677 | 51 |
| 52 | CL_T0_SG_N5 | 20160718 | 2016 | Summer | July | 0 | CL | SG | N5 | N | 5 | CL-SG | 5.16 | 4.016064257 | 51 |

*First few rows/columns shown here. See excel file for full Table S1*

| **Table S2.** Identification of variable names in found in Table S1 | | |
| --- | --- | --- |
| ID | Unit | Description |
| SampleID | NA | Identification Number of Specific Sample in the Dataset |
| SampleName | NA | Unique string of identification details for each sample |
| Date | NA | Numeric date of sample taken by year, month and day |
| Year | NA | Year sample was taken |
| Season | NA | Season of the year sample was taken |
| Month | NA | Month sample was taken |
| Months_After_SG | NA | Numeric expression of the number of months elapsed after the planting of switchgrass at each site |
| CL | NA | Clay Loam Field Site Near Ardmore, OK |
| SL | NA | Silt Loam Field Site Near Buryneville, OK |
| SG | NA | Switchgrass Plot |
| FL | NA | Fallow mixed grasses plot |
| CollarPosition | NA | Trace Gas Collar Position Location |
| Direction | Cardinal direction | Direction collar was positioned at in the field |
| Ring | NA | Position of collar from the center with 1 indicated the collar is closest to the origin |
| Subplot | NA | Site-Plot |
| pH | 0-14 | Soil pH taken by mixing 10g of soil with 50ml of ddH20, mixing and measuring in the lab via pH probe |

*First few rows shown here. See excel file for the full table*

| **Table S3.** GHG flux and community metrics differences by Wilcox Sign Ranked Test and Mann-Whitney U test for effect size between switchgrass (SG) and fallow (FL) plots. | | | | | | | | |
| --- | --- | --- | --- | --- | --- | --- | --- | --- |
| Variable | SL (SG v FL) | | | Effect Size Category | CL (SG v FL) | | | Effect Size Category |
|  | z-value | p-value | r-Effect Size |  | z-value | p-value | r-Effect Size |  |
| CO_2_ | 0.26 | 0.7932 | 0.00 | - | 3.47 | **0.000528***** | 0.13 | Small |
| CH_4_ | 9.77 | **<2.2e-16***** | 0.37 | Medium | 4.01 | **5.957e-5***** | 0.15 | Small |
| N_2_O | -0.36 | 0.7176 | 0.00 | - | 1.55 | 0.05075 | 0.00 | - |
| Observed Species | 6.60 | **4.194e-11***** | 0.38 | Medium | -1.33 | 0.184 | 0.00 | - |
| Shannon | 5.67 | **1.446e-08***** | 0.33 | Medium | -4.19 | **2.76e-05***** | 0.24 | Small |
| Simpson | 3.28 | **0.001054**** | 0.19 | Small | -5.89 | **3.767e-09***** | 0.34 | Medium |
| Chao1 | 6.04 | **1.587e-09***** | 0.35 | Medium | -0.68 | 0.49 | 0.00 | - |
| PD whole Tree | 2.72 | **0.006436**** | 0.16 | Small | -3.22 | **0.001301**** | 0.19 | Small |
| Effect size determined by Mann-Whitney U test with small (0.1 - < 0.3), medium (0.3 - < 0.5), and large (> 0.5) ranges. | | | | | | | | |

| **Table S4.** PERMANANOVA for 16S beta community structure by different treatment effects based off 999 permutations. | | | | |
| --- | --- | --- | --- | --- |
| **Group** | **PERMANOVA Table** | | | |
|  | **MeanSqs** | **F.Model** | **R^2^** | **p** |
| Time | 0.0083 | 9.248 | 0.00532 | **0.002**** |
| Site Effect (CL or SL) | 0.1152 | 128.3 | 0.07381 | **0.001***** |
| Plant Cover (SG or FL) | 0.0115 | 12.78 | 0.00735 | **0.001***** |
| Time: Plant Cover | 0.0048 | 5.377 | 0.00309 | 0.02 |
| Plant Cover: Site Effect | 0.0018 | 1.987 | 0.00114 | 0.15 |
| Time: Site Effect | 0.0911 | 101.5 | 0.05839 | **0.001***** |
| Plant Cover: Time: Site Effect | 0.0526 | 58.58 | 0.03371 | **0.001***** |

| **Table S5.** p-values from t-test between relative abundances of major phyla for SG vs FL groups of each soil type by time point. p values adjusted by conservative Bonferroni correction method. | | | | | | | | | | | | | | |
| --- | --- | --- | --- | --- | --- | --- | --- | --- | --- | --- | --- | --- | --- | --- |
| Phylum | T0 - July 2016 | | T2 - September 2016 | | T4 - November 2016 | | T8 - March 2017 | | T11 - June 2017 | | T14 - September 2017 | | T16 - November 2017 | |
|  | SL | CL | SL | CL | SL | CL | SL | CL | SL | CL | SL | CL | SL | CL |
| Acidobacteria | 0.23 | 0.19 | 0.26 | 0.14 | 0.73 | 0.2 | 0.07 | 0.61 | 0.14 | 0.51 | **<0.001***** | 0.052 | **<0.001***** | 0.99 |
| Actinobacteria | **<0.001***** | 0.28 | 0.68 | 0.095 | 0.55 | 0.44 | 0.58 | **<0.001***** | 0.95 | 0.22 | 0.5 | **<0.001***** | 0.83 | 0.65 |
| Bacteroidetes | 0.14 | 0.69 | 0.17 | **<0.001***** | 0.62 | 0.092 | 0.62 | **<0.001***** | 0.27 | 0.092 | **<0.001***** | 0.11 | **<0.001***** | 0.93 |
| Chlamydiae | 0.14 | 0.078 | 0.28 | 0.18 | 0.15 | **0.045*** | 0.65 | 0.85 | 0.97 | **<0.001***** | 0.52 | 0.7 | 0.078 | 0.336 |
| Chloroflexi | 0.056 | 0.84 | 0.49 | 0.087 | 0.13 | 0.45 | 0.14 | **0.02*** | 0.23 | 0.64 | 0.18 | **0.042*** | 0.2 | 0.63 |
| Cyanobacteria/Chloroplast | 0.56 | **<0.001***** | 0.24 | 0.071 | 0.53 | **<0.001***** | 0.051 | 0.99 | **0.034*** | 0.064 | 0.05 | **0.032*** | 0.2 | **<0.001***** |
| Deinococcus-Thermus | **<0.001***** | 0.35 | 0.12 | 0.77 | 0.92 | 0.13 | 0.89 | **<0.001***** | 0.087 | 0.075 | 0.75 | **0.015*** | **<0.001***** | 0.55 |
| Firmicutes | 0.53 | **<0.001***** | 0.34 | 0.14 | 0.36 | 0.09 | 0.42 | **<0.001***** | 0.53 | 0.76 | **<0.001***** | 0.071 | 0.63 | **0.028*** |
| Gemmatimonadetes | 0.14 | 0.18 | 0.43 | 0.98 | 0.6 | 0.52 | 0.87 | 0.068 | 0.14 | 0.069 | 0.63 | 0.37 | 0.45 | 0.064 |
| Parcubacteria | 0.49 | 0.54 | 0.066 | 0.24 | 0.36 | 0.087 | 0.094 | 0.69 | 0.34 | 0.29 | 0.38 | 0.42 | 0.36 | 0.17 |
| Plactomycetes | 0.28 | 0.54 | **<0.001***** | 0.42 | 0.25 | 0.45 | **<0.001***** | **0.021*** | 0.27 | 0.053 | 0.43 | 0.48 | 0.3 | 0.14 |
| Proteobacteria | 0.14 | 0.8 | 0.14 | 0.26 | 0.19 | 0.22 | **<0.001***** | 0.29 | 0.16 | 0.62 | 0.31 | 0.71 | 0.06 | 0.59 |
| Verrucomicrobia | 0.12 | 0.053 | 0.06 | 0.38 | 0.42 | 0.25 | 0.28 | **<0.001***** | **<0.001***** | 0.25 | 0.34 | 0.46 | **0.042*** | 0.85 |

**Supplementary Figure Text**

**Fig. S1,** **Layout for the field sites, collar positions, and soil sampling details:** **a**, layout of the SL field site and location of trace gas collars for each plot; **b**, layout of the CL field site and location of the trace gas collars for each plot; **c**, diagram of trace gas collar and chamber dimensions; **d**, diagram of collar position in relation to switchgrass plants and soil sampling details. Blue X’s represent switchgrass plants and the gray circle represents the trace gas collar.

**Fig. S2,** **Root biomass estimates and methods for:** **a**, difference between fallow and switchgrass plots for estimated root biomass by depths; **b**, graphic for how switchgrass plants were selected for root biomass estimation; **c**, graphic for how fallow root biomass estimation was conducted using four 1 m^2^ subplots in each quadrant of the plot.

**Fig. S3, Dissimilarity between soil chemistry samples through the seasons at each site between SG and FL plots for:** **a**, dissimilarity in soil carbon samples; **b**, dissimilarity in soil nitrogen content; **c**, dissimilarity in plant available phosphate content; **d**, dissimilarity in nitrate content; **e**, dissimilarity in ammonium. Ws is the slope of each line and the error associated with each slope while p-values represent the significance of each trend line. Each time point is comprised of twenty-one replicates per plot.

**Fig. S4,** **Changes in soil chemistry through the seasons at each site between SG and FL plots for:** **a**, concentration of nitrate content; **b**, ammonium content. Ws is the slope of each line and the error associated with each slope while p-values represent the significance of each trend line. Each time point is comprised of twenty-one replicates per plot.

**Fig. S5,** **Change in microbial alpha diversity measures through the seasons at each site between SG and FL plots for:** **a**, chao1 index; **b**, Shannon index. Ws is the slope of each line and the error associated with each slope while p-values represent the significance of each trend line. Each time point is comprised of twenty-one replicates per plot.

**Fig. S6,** **Correlation plot between environmental, microbial, and soil geochemistry variables**. Blue circles indicate significant positive correlations while red circles indicate negative correlations. Larger darker circles indicate a more significant and stronger correlation between variables. Positive correlations for site represent SL site while negative correspond to the CL site. ‘Microbe.Biomass’ was estimated using DNA concentrations after soil extractions. ‘observed.species’ is the total species richness (alpha diversity) of the samples. ‘PD’ represents the whole tree phylogenetic diversity of the plots. Positive correlations for ‘Plant.cover’ represent switchgrass plots and negative for fallow plots.

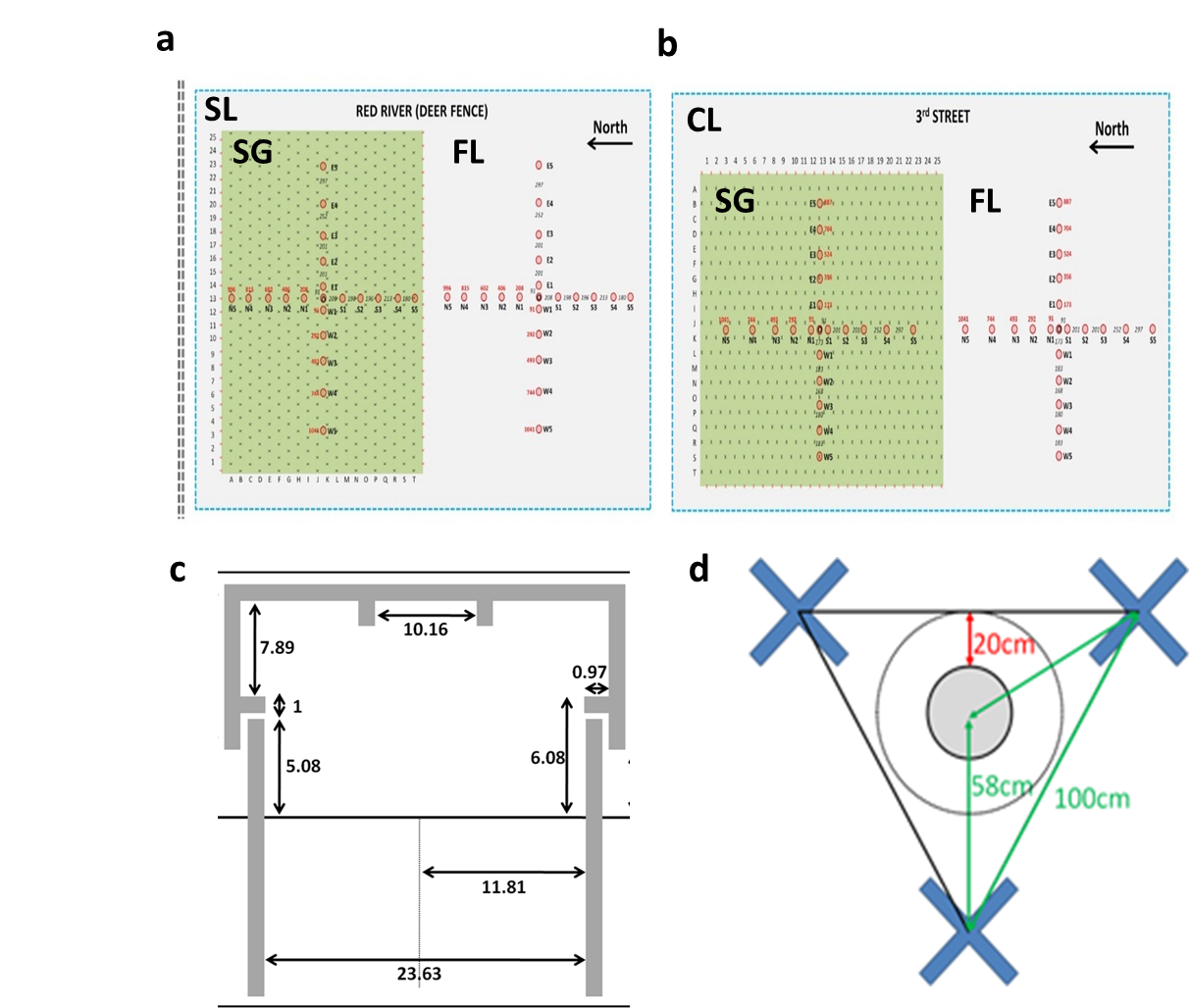

**Fig. S1**

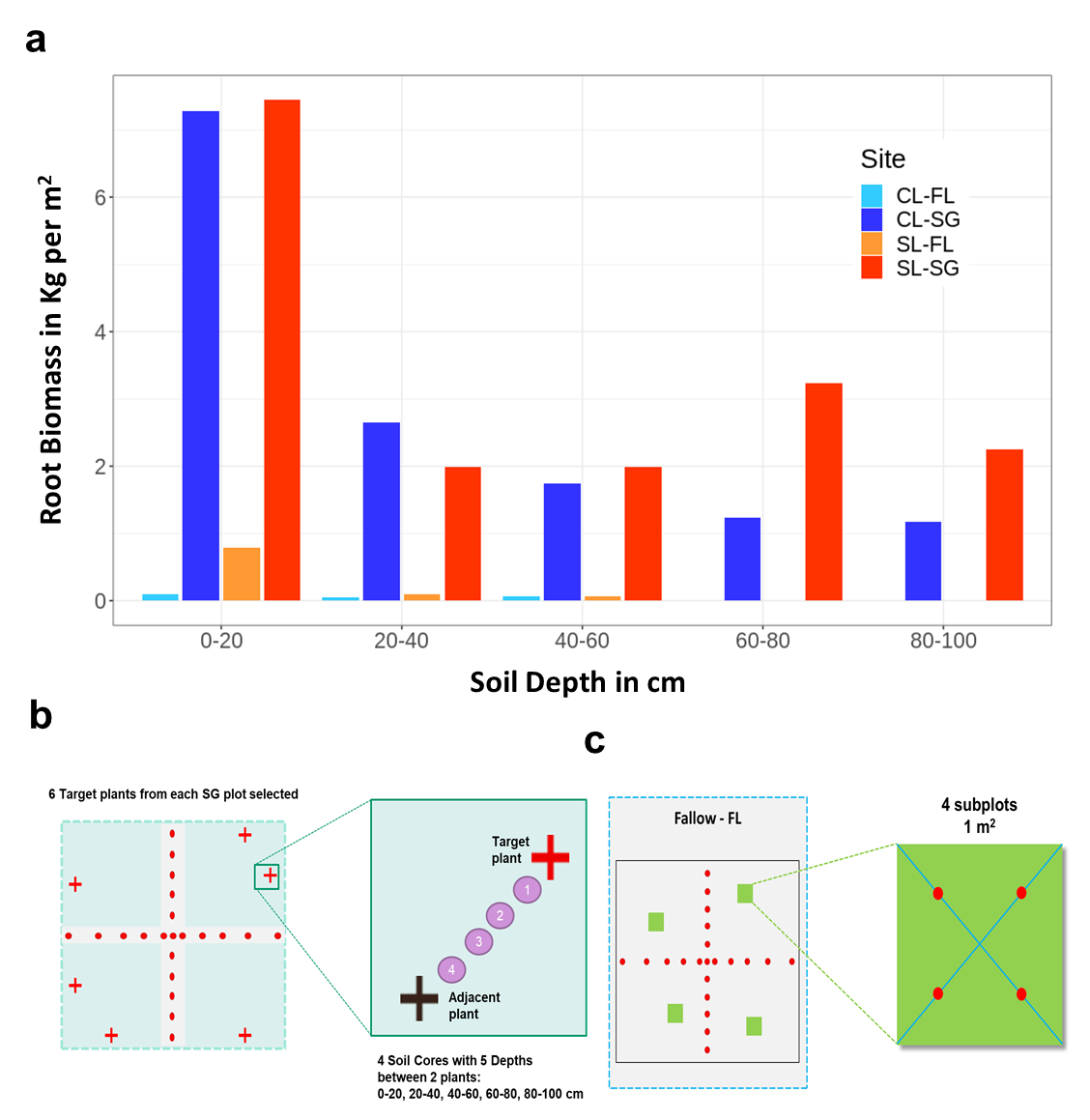

**Fig. S2**

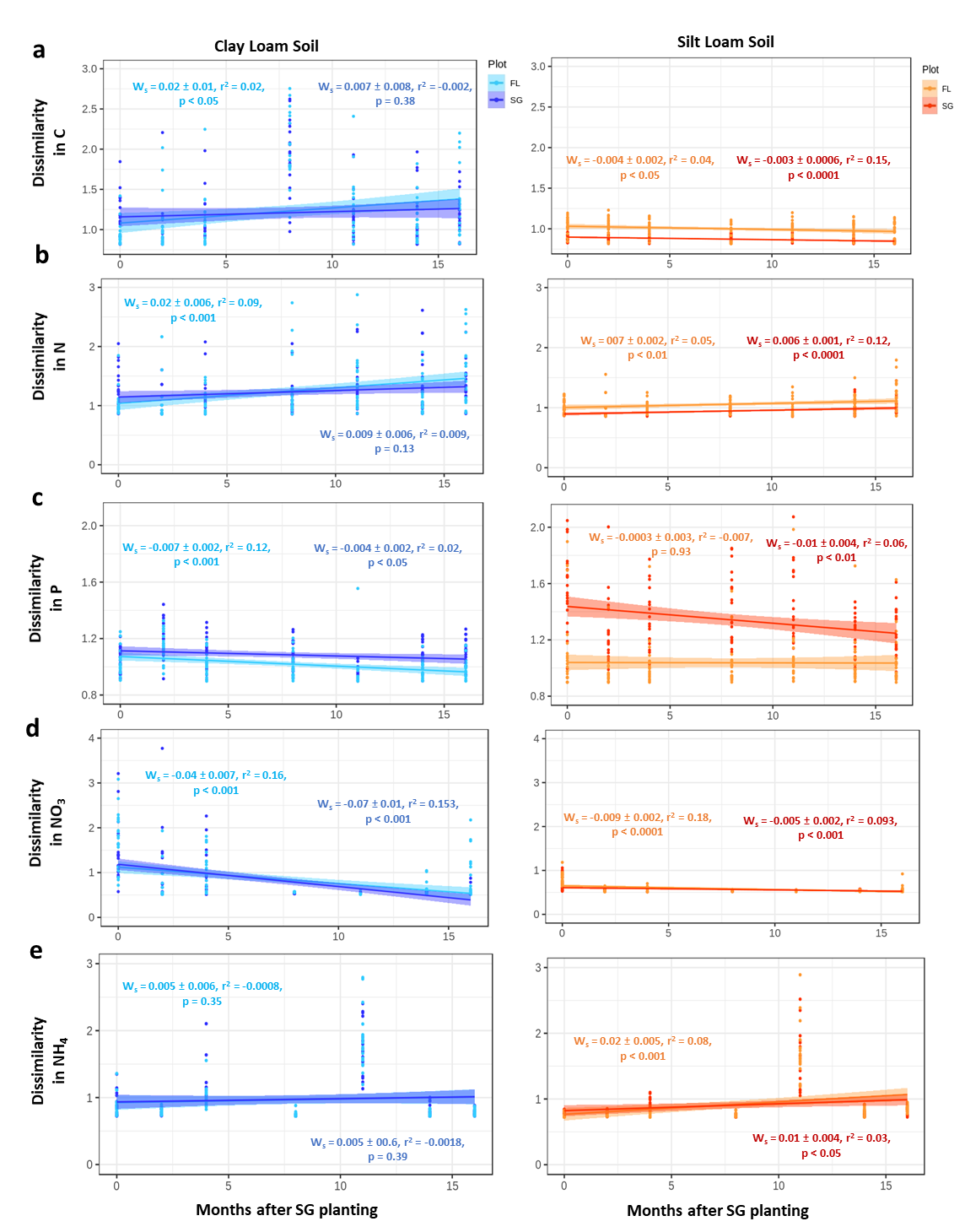

**Fig. S3**

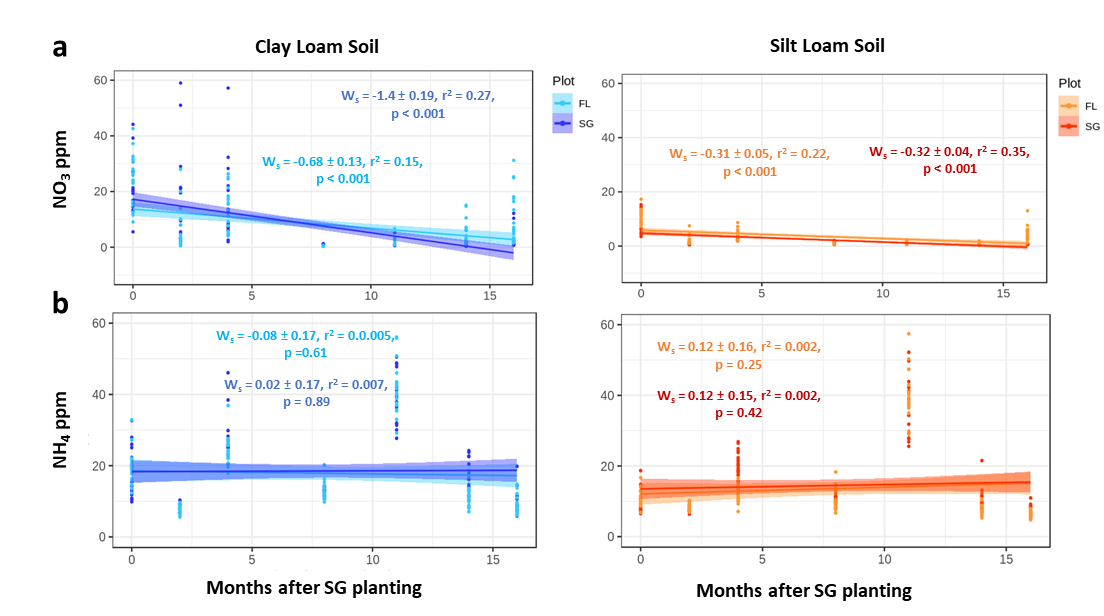

**Fig. S4**

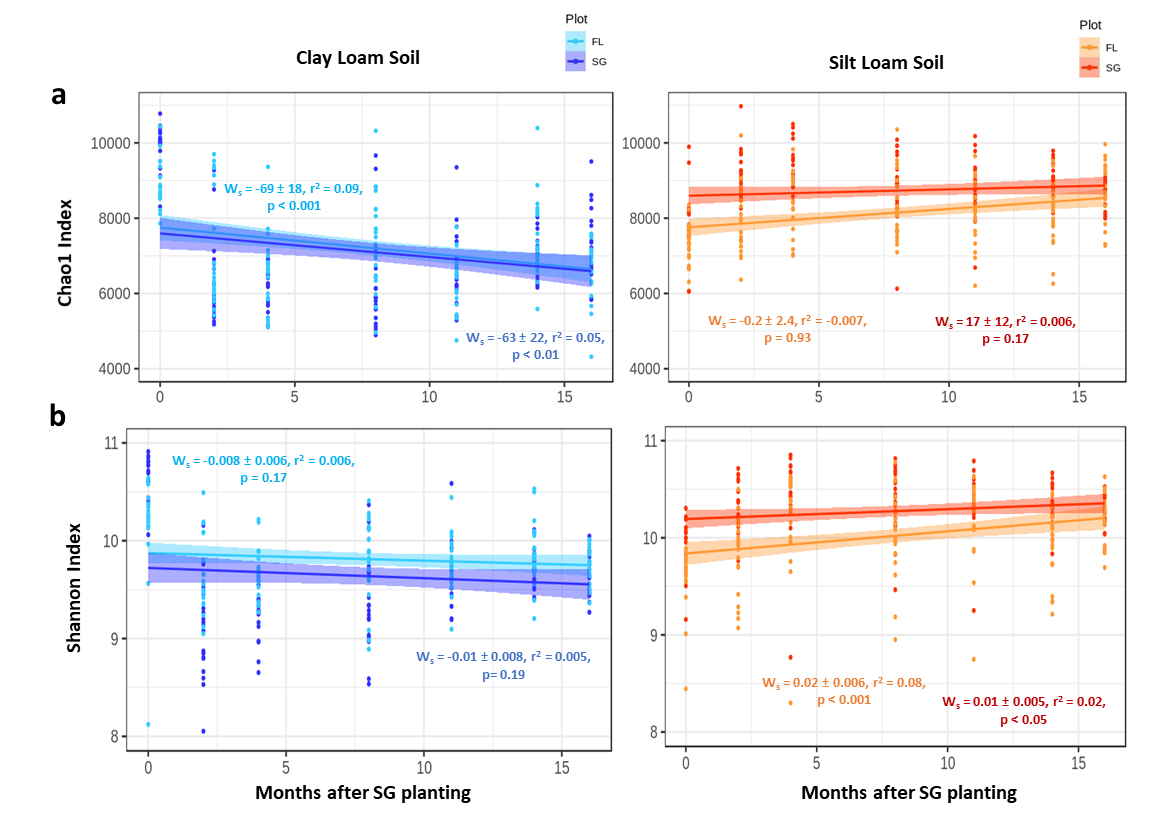

**Fig. S5**

**Supplementary Figure 3.** Change in microbial alpha diversity measures through the seasons at each site between SG and FL plots for: a, chao1 index; b, Shannon index. W_s_ is the slope of each line and the error associated with each slope while P values represent the significance of each trend line. Each time point is comprised of twenty-one replicates per plot.

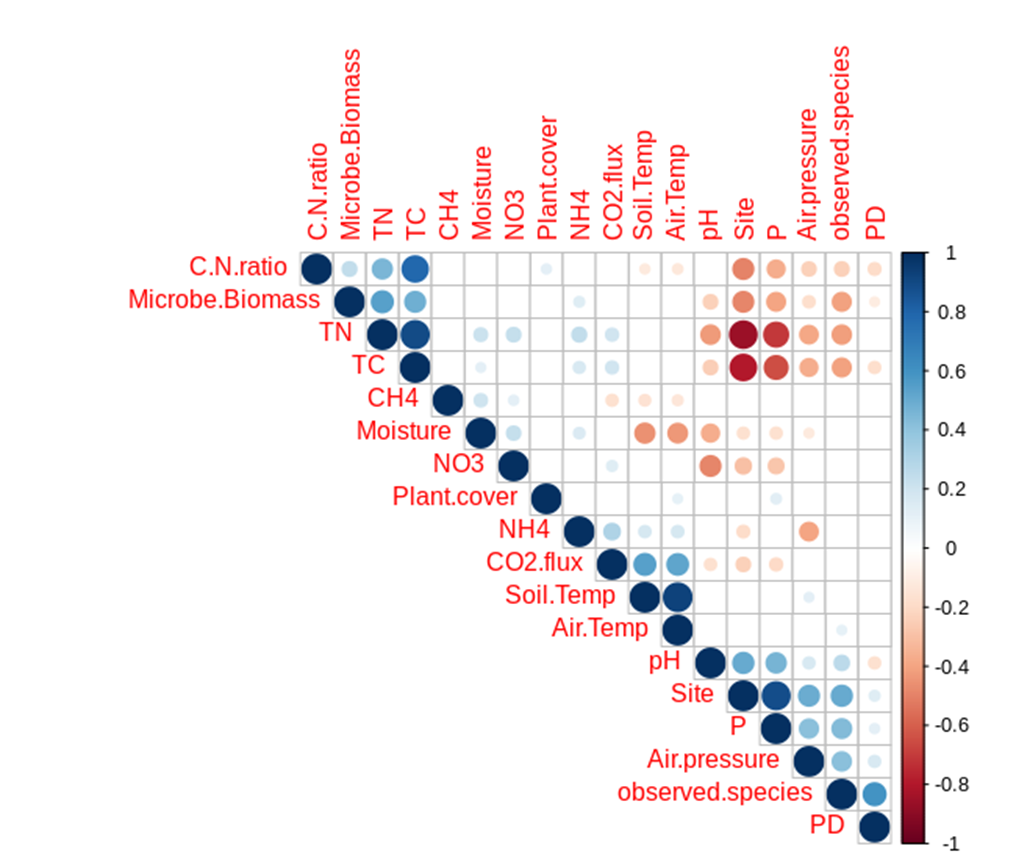

**Fig. S6**
